# PhyloRender: an interactive and agent-ready package for reproducible phylogenetic visualization

**DOI:** 10.64898/2026.11.25.769218

**Authors:** Jian He, Jingjing Chen, Zihao He, Jiamin Xiao, Lei Xie

**Affiliations:** Ecological Conservation, Restoration and Resource Utilization in Forest and Wetland Key Laboratory of Sichuan Province, Sichuan Academy of Forestry, Chengdu 610081, China; School of Ecology and Nature Conservation, Beijing Forestry University, No. 35 Tsinghua East Road, Haidian District, Beijing 100083, China

**Keywords:** chronogram, cloudogram, cytonuclear discordance, Model Context Protocol, phylogenomics, tanglegram

## Abstract

Publication-quality phylogenetic figures are central to evolutionary biology and phylogenomics. Scientific workflows are now entering the AI agent era, where AI agents already automate many routine analytical and plotting tasks. Preparing phylogenetic figures, however, still requires heavy manual work due to three bottlenecks. First, existing tree viewers and plotting packages were not built for AI agents and lack both a Model Context Protocol (MCP) server and a domain agent skill for structured, verifiable editing. Second, the standard figures of a phylogenomic paper (multi-support phylograms, fossil-calibrated chronograms, cytonuclear tanglegrams, and gene-tree cloudograms with discordance pies) cannot be made in one program, forcing files across a dozen separate tools. Third, raw plots suffer from a “last-mile” problem: support values overlap short branches, while clade grouping, nested labels, branch coloring, and geologic backgrounds take hours of manual cleanup in vector editors. We present PhyloRender, an open-source package that addresses all three bottlenecks by combining a browser workbench, a built-in MCP server, and a domain Agent Skill on one rendering engine across all four figure types. It places crowded node labels with leader lines, thickens full-support branches, aligns multi-tier clade brackets, draws International Commission on Stratigraphy (ICS) timescales, and splits tall trees into two panels without altering input topologies, branch lengths, supports, ages, or taxon names. Demonstrated on published Sapindaceae phylogenomic datasets (up to 176 taxa and 86 gene trees), PhyloRender generated publication-ready figures across all four modes with zero label overlaps (https://github.com/Jhe1004/PhyloRender).

## 1 INTRODUCTION

Phylogenetic trees are the primary visual medium for presenting evolutionary relationships, divergence times, and genealogical discordance. Interactive viewers such as FigTree (Rambaut, 2018) and iTOL (Letunic & Bork, 2024), together with scriptable libraries such as ape (Paradis & Schliep, 2019), ETE3 (Huerta-Cepas et al., 2016), and ggtree (Yu et al., 2017), have made basic tree drawing routine. At the same time, scientific computing is rapidly moving into the era of AI agents, and many multi-step data processing and plotting chores can already be carried out automatically and accurately by large language model (LLM) agents (Gao et al., 2024; Tang, 2025). Turning raw phylogenetic outputs into submission-ready journal figures has not kept pace with this shift. In practice, researchers still spend days finishing tree figures by hand because of three recurring bottlenecks.

First, existing tree software is poorly suited to AI agent workflows. When researchers ask an LLM coding or research agent to prepare a tree figure, interactive graphical user interface (GUI) viewers are inaccessible because they depend on mouse clicks, whereas general plotting packages force the agent to write custom R or Python scripts from scratch without any structured way to check whether text overlaps on the page. Current phylogenetic tools offer neither a dedicated Model Context Protocol (MCP; Anthropic, 2024) server nor a reusable domain-specific Agent Skill. Without those interfaces, an AI agent cannot reliably read raw taxon names, define clades from tip sets via their most recent common ancestor (MRCA), track figure versions, or check layout quality before exporting a file.

Second, the core evolutionary figures required in a typical phylogenomic paper are scattered across too many separate programs. A single empirical study commonly includes a main phylogram that combines support values from multiple methods such as concatenated maximum-likelihood (ML) bootstrap (BS) percentages (Minh et al., 2020) and coalescent local posterior probabilities (LPP; Zhang et al., 2018), a fossil-calibrated chronogram showing 95% highest posterior density (HPD) bars against geologic time (Douglas et al., 2022), a cytonuclear tanglegram comparing plastid or mitochondrial trees against nuclear trees, and a gene-tree conflict figure pairing a multi-locus cloudogram (Bouckaert, 2010) with four-category gene-tree discordance pie charts (Smith et al., 2015). Because no single conventional tool covers all four figure types, researchers regularly pass intermediate files across a dozen or more standalone viewers, R and Python scripts, Java command-line programs, tree-scaling utilities, and format converters. This fragmented process takes substantial time and makes it hard to keep fonts, line weights, and clade colors consistent across figures in the same manuscript.

Finally, even after initial plots are generated, a time-consuming “last-mile” editing problem remains. In rapid radiations and other regions with short internal branches, raw outputs routinely print support numbers or node ages directly over branch lines and neighboring labels. Other standard elements of a published figure also require manual work, including grouping lineages, adding nested clade names (such as tribes within subfamilies), coloring branches, and drawing shaded geologic timescale bands behind chronograms. Most researchers therefore export draft SVG or PDF files into Adobe Illustrator, CorelDRAW, or Inkscape to drag support numbers into open space, draw leader lines, line up clade brackets, and color geologic intervals by hand. That manual finishing can take hours per figure and cannot be reproduced automatically. Whenever a sample is added, an alignment is updated, or a reviewer asks for a change, the manual cleanup has to be repeated from the beginning.

Here we introduce PhyloRender, an open-source phylogenetic visualization package developed to address these three limitations for both human researchers and AI agents. PhyloRender uses a single rendering engine behind both an interactive browser workbench and a built-in MCP server paired with a domain Agent Skill, covering all four phylogenomic figure types and generating publication-ready figures from Newick and Nexus tree files without manual post-processing in external vector editors.

## 2 DESIGN AND IMPLEMENTATION

### 2.1 Browser Workbench and AI Agent Interface

PhyloRender is built around a single shared rendering engine that serves both human users and AI agents (Figure 1). Whether a researcher interacts with the browser workbench or an AI agent invokes tools through MCP, tree parsing, data validation, label placement, and vector or raster export all run through the exact same layout engine on a unified page coordinate system. In the browser workbench, users can reroot trees, color lineages, define nested clades, and adjust typography and spacing directly on the canvas.

**FIGURE 1.**
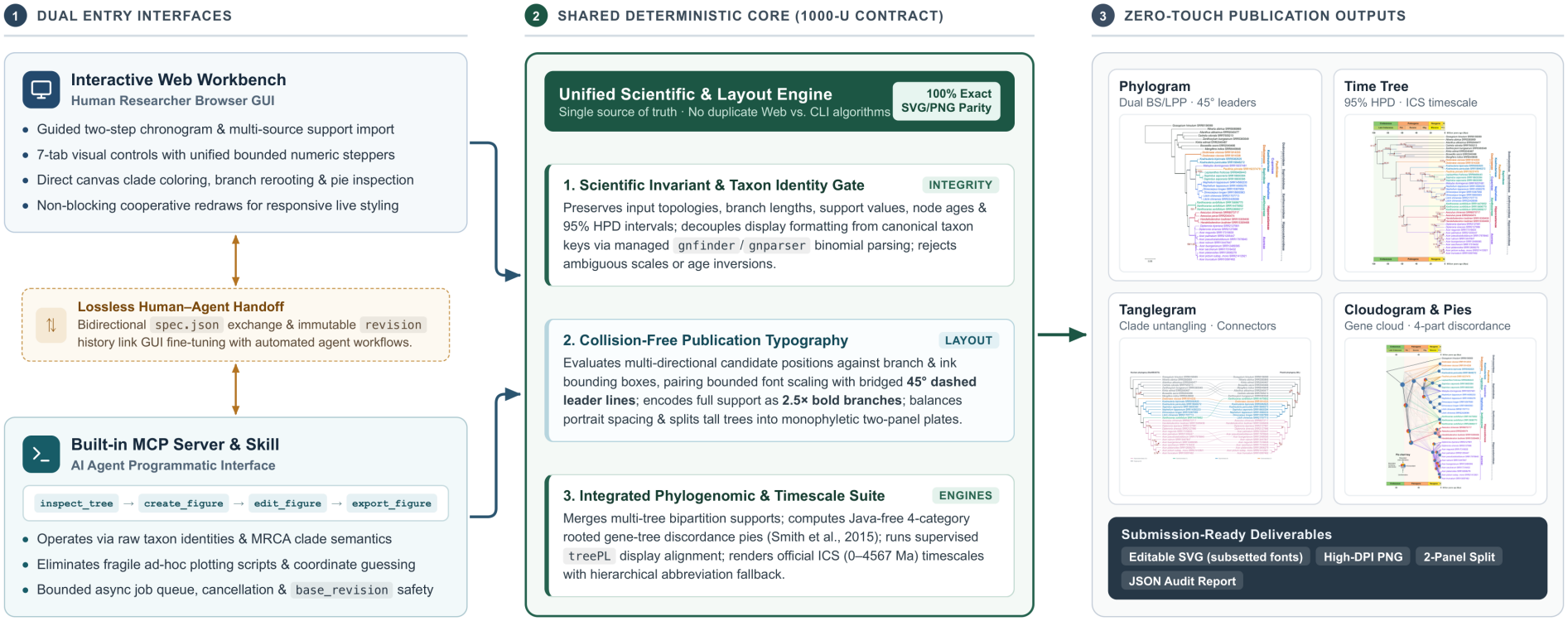
Overview of the PhyloRender workflow for human users and AI agents. Both the browser workbench and the built-in MCP server paired with the Agent Skill operate on a single shared rendering engine, exchanging figure configurations and versioned revisions across all four workspaces to export vector SVG and high-DPI PNG figures.

For AI-assisted workflows, the built-in MCP server exposes structured tools covering the full plotting cycle: agents can inspect original tip names and taxon overlap across input trees, create or update figures through versioned revisions, define clades by MRCA tip sets, and verify layout quality before exporting finished graphics. Alongside the server, a companion Agent Skill defines domain-specific plotting rules, such as confirming outgroups with the user before rerooting an undated tree and reporting taxon mismatches before constructing a tanglegram (runtime architecture in Supplementary Methods 1; user manual, configurable options, and operational workflows in Supplementary Methods 2). PhyloRender is not an autonomous agent itself; the MCP server supplies the programmatic interface while the skill encodes practical phylogenomic plotting conventions. Because both the browser workbench and the MCP server read and write the same figure configuration, and because each programmatic update returns a structured report verifying topology preservation and zero label overlap, human researchers and AI agents can transfer figures between interactive and automated sessions without losing styling or revision history.

### 2.2 Four Core Phylogenetic Figure Types

PhyloRender integrates the four standard phylogenomic figure types within a single environment sharing consistent typography and color styling. In the Phylogram workspace, a main tree can be paired with additional support trees from other inference methods; corresponding branches are matched automatically across trees even when rooting differs, so ML bootstrap percentages and coalescent or Bayesian posterior probabilities can be combined on one topology without manual node mapping. In the Time Tree workspace, fossil-calibrated chronograms preserve their supplied root and divergence ages, omit node support values so chronological intervals remain unambiguous, and display 95% HPD bars against an official ICS geologic timescale (Cohen et al., 2013). In the Tanglegram workspace, two conflicting phylogenies such as nuclear and organellar trees are placed side by side, original tip identifiers are matched one to one, and sister clades are rotated to minimize crossing connectors between shared taxa without altering either tree’s topology.

For multi-locus conflict across gene-tree sets, the Cloudogram workspace combines a dated species tree with a collection of ML gene trees in one view to display topological discordance alongside four-category node pie charts on a shared geologic timescale. Conventionally, assembling such a figure requires installing two external command-line tools: the Java program PhyParts (Smith et al., 2015) to count gene-tree conflict and the C++ program treePL (Smith & O’Meara, 2012), together with its external C++ mathematical libraries, to scale gene-tree branches onto the species-tree timescale. To make this workflow self-contained without external software installations, PhyloRender implements both supporting calculations directly in JavaScript (algorithmic details in Supplementary Methods 1). A built-in discordance module applies the clade comparison rules of Smith et al. (2015) to count concordant, dominant alternative, remaining alternative, and uninformative gene trees at each species-tree node, including gene trees with missing taxa, while keeping all four counts independent of input tree order. A companion branch-scaling module implements the rate-smoothed penalized-likelihood method of Sanderson (2002) and Smith and O’Meara (2012) while holding the ages of clades shared with the dated species tree fixed, scaling the remaining gene-tree branches smoothly onto the species-tree timescale without numerical failures on zero-length or very short branches. Output metadata explicitly record that these scaled gene-tree branch lengths serve solely for visual alignment rather than independent divergence-time estimation.

### 2.3 Automatic Layout and Data Integrity

In trees with dense branching or short internal edges, raw plots often print node numbers directly over branches and neighboring text, requiring manual cleanup in external vector graphics editors. PhyloRender avoids this post-processing step by placing crowded node annotations automatically during layout. For each displayed support value or node age, the engine evaluates candidate positions around the node against all branch lines, node markers, tip names, and neighboring numbers. Where short internal branches leave no open space beside a split, the label is moved into nearby whitespace and connected back to its node by a leader line (placement algorithm in Supplementary Methods 1). Along well-supported backbones, branches reaching full support across all matched trees can be drawn with thickened lines rather than repeating 100/1.00 labels, whereas values below a chosen cutoff are masked as –.

Clade brackets, geologic timescales, tip names, and multi-panel pagination are formatted automatically under the same layout engine. Clades defined by MRCA tip sets can be annotated as single brackets or nested taxonomic tiers (such as tribes within subfamilies), coloring descendant lineages and aligning each taxonomic rank into a clean vertical column without overlapping labels. On chronograms and cloudograms, ICS geologic bands fit period and epoch labels cleanly within even narrow intervals, while tip-label parsing italicizes Latin binomials and keeps specimen voucher codes upright. For tall phylograms and chronograms that exceed a single page, a two-panel option splits the tree at a monophyletic clade into two equal-width panels and continues any spanning outer clade bracket across both panels.

Built-in integrity checks prevent visual styling from altering underlying phylogenetic data. Original tip identifiers are kept separate from formatted display labels and serve as the sole matching key across trees, so italicizing species names or converting separators never alters specimen voucher or accession codes. Ladderization and tanglegram untangling swap sister clades only and verify topology preservation prior to export. Chronograms preserve their input root, node ages, and HPD intervals, and support-value imports reject files whose values cannot be distinguished clearly between 0–1 probabilities and 0–100 percentages rather than guessing units.

## 3 EMPIRICAL DEMONSTRATION AND VALIDATION

We demonstrated PhyloRender across all four workspaces using published phylogenomic datasets of the soapberry family, Sapindaceae, extracted from He et al. (2025; subfamilial and tribal classification following Buerki et al., 2021), with every figure exported directly from PhyloRender without manual editing in external vector graphics software. To test node-label placement across rapid short-branch radiations, we extracted a 50-taxon Sapindoideae crown clade from the 176-taxon Sapindaceae phylogeny of He et al. (2025), retaining unmodified ML branch lengths and paired ML BS / ASTRAL LPP supports, ladderized it identically, and compared unedited outputs from the R package ape (v5.8.1), the Python toolkit ETE3 (v3.1.3), and PhyloRender (Figure 2, Table S1). Both ape (Figure 2b) and ETE3 in its branch-length-preserving floating mode (Figure 2c) stacked overlapping composite support labels along the short radiation backbone while repeating 100/1.00 strings across full-support branches. PhyloRender (Figure 2d) preserved exact branch lengths and topology, drew full-support branches with thickened lines, and placed all remaining composite labels with leader lines without overlap.

**FIGURE 2.**
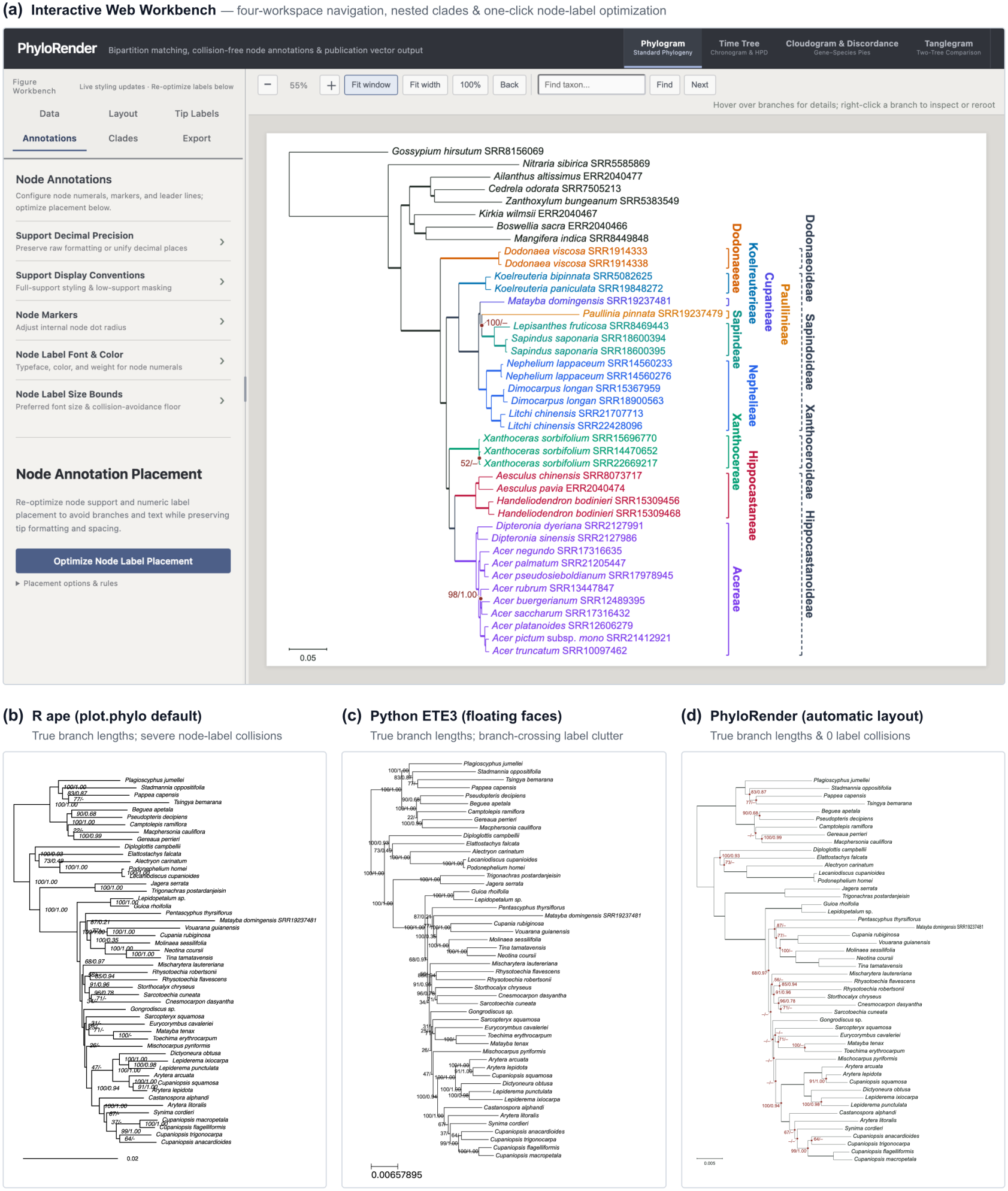
Interactive Web Workbench and node-label comparison on a 50-taxon rapid radiation. **(a)** User interface of the interactive PhyloRender Web Workbench. **(b–d)** Unedited outputs on the same ladderized 50-taxon Sapindoideae crown clade (dataset from He et al., 2025), retaining unmodified ML branch lengths and paired ML BS / ASTRAL LPP supports (invocation details in Table S1): **(b)** ape (v5.8.1) prints composite labels at fixed node offsets, producing heavy overlap along short backbone branches; **(c)** ETE3 (v3.1.3, position=“float”, preserving true branch lengths) stacks floating labels across branches and repeats 100/1.00 strings on full-support edges; **(d)** PhyloRender preserves exact branch lengths and topology, draws full-support branches with thickened lines, and places all remaining composite supports with leader lines without overlap.

Using the broader Sapindaceae datasets of He et al. (2025), we generated a coordinated set of manuscript figures under shared typography and clade colors (Figure 3, Figure 4, and Table S2). The 41-taxon concatenated ML and ASTRAL phylogram (Figure 3a) combined dual-method supports automatically and aligned two-tier tribe and subfamily brackets across all 33 Sapindaceae ingroup accessions without label overlap. The companion 41-taxon StarBEAST3 chronogram (Figure 3b) displayed divergence ages and 95% HPD bars against an official 0–100 Ma ICS geologic timescale with zero text collisions, while the cytonuclear tanglegram comparing the nuclear and plastid trees across their 29 shared accessions (Figure 3c) untangled taxon connectors via sister-clade rotation to highlight plastid–nuclear topological conflict cleanly.

**FIGURE 3.**
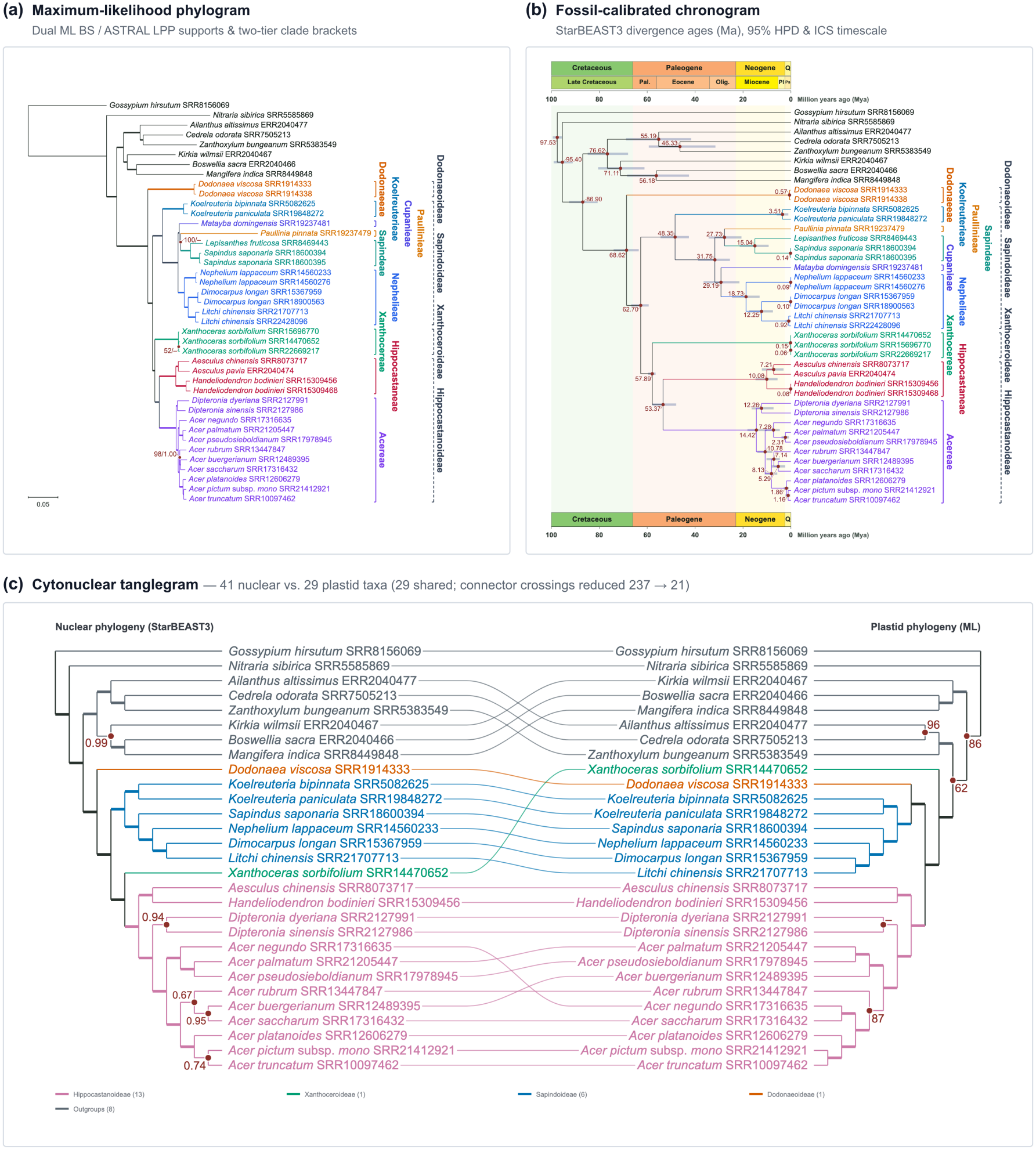
Sapindaceae case studies across the Phylogram, Time Tree, and Tanglegram workspaces (datasets from He et al., 2025). **(a)** A 41-taxon concatenated ML phylogram paired automatically with ASTRAL LPP supports and aligned twotier tribe and subfamily brackets across all 33 Sapindaceae ingroup accessions. **(b)** A 41-taxon StarBEAST3 fossil-calibrated chronogram showing internal node ages, 95% HPD bars, an official ICS geologic timescale (0–100 Ma), and aligned two-tier brackets. **(c)** A cytonuclear tanglegram comparing the 41-taxon nuclear tree with a 29-taxon plastid tree across 29 shared accessions, using sister-clade rotation to untangle taxon connectors.

**FIGURE 4.**
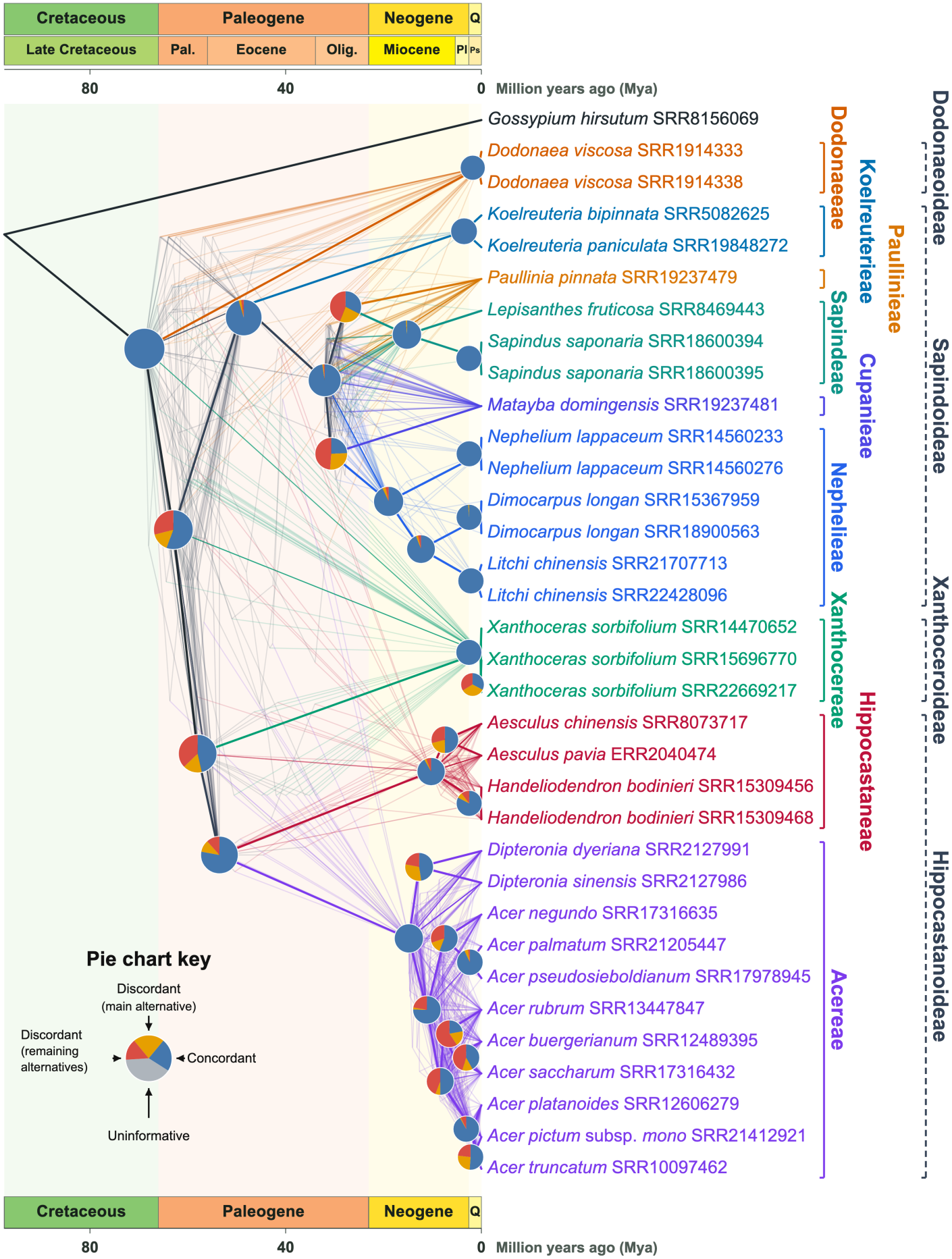
Gene-tree and species-tree cloudogram with four-category discordance pie charts for Sapindaceae (dataset from He et al., 2025). A 34-taxon dated species tree is overlaid with 86 ML gene trees scaled to shared clade ages by PhyloRender’s built-in penalized-likelihood module (following Sanderson, 2002; Smith & O’Meara, 2012) for visual display, paired with four-category gene-tree discordance pie charts (concordant, dominant alternative, remaining alternatives, uninformative) calculated by PhyloRender’s built-in module following Smith et al. (2015), official ICS geologic bands, species-tree MRCA branch coloring, and aligned vertical two-tier clade brackets.

In the Cloudogram workspace, combining a 34-taxon dated species tree with 86 ML gene trees (Figure 4) produced four-category discordance pie charts, penalized-likelihood visual gene-tree overlays, MRCA-colored gene branches, an ICS timescale, and aligned two-tier brackets in one workflow with zero overlaps. On the full 176-taxon Sapindaceae phylogeny (Figure S1), two-panel export isolated the monophyletic crown Sapindoideae clade into a second equal-width panel, carried the spanning outer subfamily bracket across both panels, and placed all composite support labels without collision.

Finally, we benchmarked both built-in JavaScript modules against reference implementations across simulated trees and empirical plant phylogenomic datasets containing up to 8,467 gene trees (Tables S3 and S4). Across seven discordance test suites covering all 720 five-taxon rooted binary and multifurcating tree combinations, multiple support cutoffs, random input-order permutations, and four empirical locus sets with complete and missing-taxon sampling, PhyloRender matched an independent Python reference implementation across all four category counts with 100% exact node agreement (maximum discrepancy 0; Table S3). Across penalized-likelihood scaling benchmarks spanning rate-shifted simulated trees, zero- and micro-branch radiations, ten orders of magnitude in root-age scale, and empirical ML gene-tree sets (17,499 evaluated internal nodes in total), PhyloRender closely matched treePL (Smith & O’Meara, 2012; *r ≥* 0.9999989 across all suites, with mean absolute error *≤* 2.6 × 10^*−*5^ Ma on well-resolved sets; Table S4) while running 26 to 138 times faster. Cross-interface verification also confirmed that the browser workbench and MCP server export identical SVG files from the same configuration.

## 4 DISCUSSION

As biological research moves into the AI agent era, sequence processing, multi-step bioinformatics pipelines, and downstream statistical analyses are increasingly orchestrated by LLM agents (Gao et al., 2024; Hayakawa & Ozaki, 2026; Tang, 2025), yet publication-grade figure preparation has remained a persistent bottleneck. Asking a general-purpose agent to write one-off R or Python plotting scripts from scratch leaves two practical gaps: the agent receives no geometric feedback on whether rendered text overlaps on crowded short branches, and ad hoc generated code lacks domain safeguards against silent changes to specimen identifiers, rooting, or support scales, a reliability challenge shared across biomedical LLM applications when models are not grounded in validated domain tools (Lobentanzer et al., 2025). PhyloRender demonstrates an alternative design pattern for scientific visualization software in this setting. By placing a shared layout and integrity-checking engine behind both a standardized MCP server (Anthropic, 2024) and a domain-specific Agent Skill, the software allows AI agents to inspect raw tree properties, follow phylogenetic plotting conventions, and verify zero label collisions programmatically, while enabling researchers to open the exact same figure configuration in the browser workbench for interactive visual refinement.

Within the existing ecosystem of tree visualization tools, established packages continue to serve specialized roles well: scriptable libraries such as ggtree (Yu et al., 2017), ape (Paradis & Schliep, 2019), and ETE3 (Huerta-Cepas et al., 2016) excel at custom multi-track data integration through code, iTOL supports web-based interactive exploration and circular ring annotations (Letunic & Bork, 2024), and DensiTree visualizes posterior tree distributions from Bayesian analyses (Bouckaert, 2010). PhyloRender complements these tools by targeting the four core phylogenetic figures needed in most empirical phylogenomic manuscripts and removing two sources of routine friction: cross-platform installation barriers for cloudogram conflict and branch-scaling calculations, and hours of manual cleanup in external vector graphics editors. In conventional workflows, exporting a draft tree into a vector editor breaks reproducibility because manual label dragging and bracket alignment cannot be replayed when taxa are added, outgroups are revised, or analyses are updated during peer review. Recording clade definitions, color schemes, and layout settings in a reusable configuration allows either a researcher or an AI agent to regenerate updated, overlap-free figures directly from revised tree files.

At present, PhyloRender focuses on these four standard phylogenomic figure types rather than arbitrary peripheral data tracks; studies requiring complex side-by-side heatmaps, gene-structure diagrams, or multi-layered ecological matrices alongside a tree will still benefit from grammar-of-graphics scripting packages such as ggtree. At the same time, PhyloRender is released under an open-source license not only as a ready-to-use application, but also as a modular framework and practical reference example for agent-accessible scientific software. Because the rendering modules, configuration schema, MCP tools, and Agent Skill are cleanly separated and openly documented, researchers can use coding AI agents directly to extend the codebase and add custom plotting features, layout styles, or specialized annotation tracks tailored to their own studies while retaining the platform’s collision-free layout and data-integrity checks.

## AUTHOR CONTRIBUTIONS

**Jian He:** Conceptualization; data curation; formal analysis; investigation; methodology; project administration; software; validation; visualization; writing – original draft; writing – review and editing. **Jingjing Chen:** Software; writing – review and editing. **Zihao He:** Writing – review and editing. **Jiamin Xiao:** Data curation; software; supervision; validation; writing – original draft; writing – review and editing. **Lei Xie:** Funding acquisition; resources; supervision; writing – review and editing.

## ACKNOWLEDGEMENTS

We thank the developers of treePL, PhyParts, gnfinder, gnparser, and the International Commission on Stratigraphy for openly documenting the algorithms, tools, and chronostratigraphic standards referenced or integrated here.

## ARTIFICIAL INTELLIGENCE DISCLOSURE

Gemini 4 Carbon (Google) was used as an assistive tool during software development and to polish the English language of the manuscript. All authors reviewed and verified the code and text, and take full responsibility for the content of this work.

## DATA ACCESSIBILITY AND BENEFIT-SHARING

### Data and Software Accessibility

PhyloRender (source code, Web Workbench, MCP server, companion Agent Skill, bilingual documentation, and validation test suites) is openly available under the GNU General Public License v3.0 or later (GPL-3.0-or-later) at https://github.com/Jhe1004/PhyloRender, and a permanent release archive will be deposited in Zenodo upon acceptance. Empirical Sapindaceae phylogenomic datasets underlying Figures 2–4 and Figure S1 were obtained from He et al. (2025; https://doi.org/10.3390/ijms26041581), and the exact demonstration subsets and figure configurations will be archived with a permanent DOI upon acceptance.

### Benefit-Sharing Statement

No physical biological specimens or genetic resources were newly collected for the development of this software. Benefits from this work are shared through the open-source release of PhyloRender, its MCP server, Agent Skill, and bilingual documentation for free public use.

## SUPPORTING INFORMATION

Additional supporting information can be found online in the Supporting Information section at the end of this article:

**Supplementary Methods 1**. Mathematical formulations of PhyloRender’s layout, untangling, discordance, and branch-scaling algorithms, together with its shared runtime and transactional consistency architecture.

**Supplementary Methods 2**. User manual, configurable options mapped across the Web Workbench, MCP server, and Agent Skill for all four workspaces, and step-by-step operational workflows.

**Figure S1.** Automated two-panel split at a monophyletic clade, aligned two-tier tribe and subfamily brackets carried across panels, and dual-source support labels on the full 176-taxon Sapindaceae phylogeny.

**Table S1**. Software versions, input files, and exact commands used for the node-label comparison in Figure 2b–d.

**Table S2**. Layout diagnostics, invariant checks, and run metrics across all main and supplementary empirical figures.

**Table S3**. Benchmark verification of PhyloRender’s built-in gene-tree discordance module across simulated and empirical datasets.

**Table S4**. Node-age agreement and runtime comparison between PhyloRender’s built-in penalized-likelihood branch-scaling module and treePL.

