## Supporting Information for "PhyloRender: an interactive and agent-ready package for reproducible phylogenetic visualization"

**Article Title:** *PhyloRender: an interactive and agent-ready package for reproducible phylogenetic visualization*
**Journal:** *Molecular Ecology Resources* (Software Note)
**Authors:** Jian He, Jingjing Chen, Zihao He, Jiamin Xiao, and Lei Xie

This index summarizes all individual Supporting Information files accompanying the manuscript and lists the literature cited in each file. Full mathematical formulations, user guides, figure legends, tables, and per-file reference lists are provided inside the respective standalone files.

#### Inventory of Supporting Information Files

| Item | Standalone File(s) | Concise Summary of Contents | References Cited in File |
|---|---|---|---|
| **Supplementary Methods 1** | `Supplementary_Methods_1.md` | Algorithmic formulations and system consistency architecture: safe Newick/Nexus parsing, raw-identity decoupling, unrooted branch matching, score-guided node-label placement and leader routing, hierarchical clade brackets, ICS timescale fitting, two-panel splitting, tanglegram sister-clade untangling, built-in gene-tree discordance and penalized-likelihood visual alignment, shared headless runtime, and transactional revision control (Sections S1.1–S1.6). | Anthropic (2024); Cohen et al. (2013); Sanderson (2002); Smith & O'Meara (2012); Smith et al. (2015); Zhang et al. (2018) |
| **Supplementary Methods 2** | `Supplementary_Methods_2.md` | User manual and operational reference: workspace selection guide, shared and per-workspace configurable options mapped across the Web Workbench, MCP server parameters/edit operations, and Agent Skill natural-language prompts, inventory of all 15 MCP tools and five domain confirmation rules, and step-by-step human and AI agent workflows (Sections S2.1–S2.5). | Anthropic (2024); Cohen et al. (2013); Sanderson (2002); Smith & O'Meara (2012); Smith et al. (2015); Zhang et al. (2018) |
| **Figure S1** | `Figure_S1.md` and `figS1_sapindaceae_176tips_panels/FigS1_sapindaceae_176tips_panels.pdf` (`.png`) | Automated monophyletic two-panel split, cross-panel two-tier tribe and subfamily brackets, and dual-method support labels on the full 176-taxon Sapindaceae phylogeny. | Buerki et al. (2021); He et al. (2025); Zhang et al. (2018) |

| Item | Standalone File(s) | Concise Summary of Contents | References Cited in File |
|---|---|---|---|
| **Table S1** | `Table_S1.md` | Software versions, input files, and rendering commands used for the 50-taxon Sapindoideae node-label comparison in Figure 2b–d. | He et al. (2025); Huerta-Cepas et al. (2016); Paradis & Schliep (2019) |
| **Table S2** | `Table_S2.md` | Empirical dataset summary, displayed annotations, and automated layout and data-integrity verification across Figures 2–4 and Figure S1. | Buerki et al. (2021); Cohen et al. (2013); Douglas et al. (2022); He et al. (2025); Sanderson (2002); Smith & O'Meara (2012); Smith et al. (2015) |
| **Table S3** | `Table_S3.md` | Numerical equivalence benchmarks of PhyloRender's built-in JavaScript gene-tree discordance module across simulated and empirical phylogenomic datasets. | He et al. (2025); Smith et al. (2015) |
| **Table S4** | `Table_S4.md` | Node-age agreement and runtime comparison between PhyloRender's built-in JavaScript penalized-likelihood branch-scaling module and `treePL`. | He et al. (2025); Sanderson (2002); Smith & O'Meara (2012) |

#### Consolidated References Cited Across Supporting Information

Anthropic. (2024). *Model Context Protocol specification*. https://modelcontextprotocol.io/

Buerki, S., Callmander, M. W., Acevedo-Rodríguez, P., Lowry, P. P., II, Munzinger, J., Bailey, P., Maurin, O., Brewer, G. E., Epitawalage, N., Baker, W. J., & Forest, F. (2021). An updated infra-familial classification of Sapindaceae based on targeted enrichment data. *American Journal of Botany*, *108*(7), 1234–1251. https://doi.org/10.1002/ajb2.1693

Cohen, K. M., Finney, S. C., Gibbard, P. L., & Fan, J.-X. (2013). The ICS International Chronostratigraphic Chart. *Episodes*, *36*(3), 199–204. https://doi.org/10.18814/epiiugs/2013/v36i3/002

Douglas, J., Jiménez-Silva, C. L., & Bouckaert, R. (2022). StarBEAST3: Adaptive parallelized Bayesian inference under the multispecies coalescent. *Systematic Biology*, *71*(4), 901–916. https://doi.org/10.1093/sysbio/syac010

He, J., Li, M., Wu, H., Cheng, J., & Xie, L. (2025). Unraveling the ancient introgression history of *Xanthoceras* (Sapindaceae): Insights from phylogenomic analysis. *International Journal of Molecular Sciences*, *26*(4), 1581. https://doi.org/10.3390/ijms26041581

Huerta-Cepas, J., Serra, F., & Bork, P. (2016). ETE 3: Reconstruction, analysis, and visualization of phylogenomic data. *Molecular Biology and Evolution*, *33*(6), 1635–1638. https://doi.org/10.1093/molbev/msw046

Paradis, E., & Schliep, K. (2019). ape 5.0: An environment for modern phylogenetics and evolutionary analyses in R. *Bioinformatics*, *35*(3), 526–528. https://doi.org/10.1093/bioinformatics/bty633

Sanderson, M. J. (2002). Estimating absolute rates of molecular evolution and divergence times: A penalized likelihood approach. *Molecular Biology and Evolution*, *19*(1), 101–109. https://doi.org/10.1093/oxfordjournals.molbev.a003974

Smith, S. A., Moore, M. J., Brown, J. W., & Yang, Y. (2015). Analysis of phylogenomic datasets reveals conflict, concordance, and gene duplications with examples from animals and plants. *BMC Evolutionary Biology*, *15*, 150. https://doi.org/10.1186/s12862-015-0423-0

Smith, S. A., & O'Meara, B. C. (2012). treePL: Divergence time estimation using penalized likelihood for large phylogenies. *Bioinformatics*, *28*(20), 2689–2690. https://doi.org/10.1093/bioinformatics/bts492

Zhang, C., Rabiee, M., Sayyari, E., & Mirarab, S. (2018). ASTRAL-III: Polynomial time species tree reconstruction from partially resolved gene trees. *BMC Bioinformatics*, *19*(Suppl 6), 153. https://doi.org/10.1186/s12859-018-2129-y

### Supplementary Methods 1: Algorithmic Formulations and System Consistency Architecture

**Article Title:** *PhyloRender: an interactive and agent-ready package for reproducible phylogenetic visualization*
**Journal:** *Molecular Ecology Resources* (Software Note)
**Authors:** Jian He, Jingjing Chen, Zihao He, Jiamin Xiao, and Lei Xie

This file documents the mathematical formulations, layout geometry, built-in phylogenetic conflict and branch-scaling solvers, and runtime consistency architecture underlying PhyloRender. For user-oriented operational workflows and a complete reference of configurable options across the Web Workbench, MCP server, and Agent Skill, see Supplementary Methods 2.

#### S1.1 | Tree Parsing, Taxon Identity Decoupling, and Unrooted Branch Matching

##### Safe Newick and Nexus parsing

PhyloRender parses standard Newick and Nexus tree files through a deterministic single-pass tokenizer that preserves raw leaf identifiers, branch lengths (rejecting negative or non-numeric values rather than zeroing them silently), internal node labels, and square-bracketed metadata comments (`[&...]`) such as Bayesian divergence times and 95% highest posterior density (HPD) intervals. For Nexus inputs, single- and double-quoted strings and nested square-bracket comment depth are tracked simultaneously while reading strictly within the boundaries of the `TREES` block; consequently, numbers or text inside annotation blocks or other Nexus blocks are never mistaken for `TRANSLATE` taxon mappings. Multi-tree Nexus files or concatenated multi-tree Newick inputs supplied to single-tree workspaces are rejected with an explicit blocking error instructing the user or caller to select or summarize a single consensus or maximum clade credibility (MCC) tree externally rather than defaulting silently to the first posterior sample.

##### Decoupling raw taxon identities from formatted display names

Across all workspaces, every terminal node retains two separate representations: an immutable raw taxon identifier extracted directly from the input tree file and a display label used solely for canvas rendering. Cross-tree taxon matching, outgroup rooting, most recent common ancestor (MRCA) clade definitions, and topology verification always operate on the raw identifiers. When constructing default display labels, PhyloRender splits each raw label immediately before the first token containing digits (corresponding to a specimen voucher or sequence accession code): underscores in the leading taxonomic name prefix are converted to spaces, whereas trailing specimen or accession tokens containing internal underscores (such as `2023080906_1` or `SRR21412921`) are kept upright and unmodified. When automatic taxonomic name formatting is active, Latin genus names and infrageneric, specific, and infraspecific epithets are italicized, while taxonomic rank, open-nomenclature identification, and hybrid qualifiers (`subsp.`, `ssp.`, `var.`, `subvar.`, `f.`, `forma`, `sp.`, `cf.`, `aff.`, `sect.`, `ser.`, and hybrid `×`) are kept upright; when rank standardization is enabled (on by default), variant rank spellings such as `ssp.`/`subspecies`, `forma`/`fo.`, and `variety` are normalized in roman type to `subsp.`, `f.`, and `var.`, respectively. Custom display-name mapping tables (CSV, TSV, TXT, or `.xlsx` files) validate that every source key resolves uniquely to an existing tip on the active tree without conflicting duplicate rows (multiple distinct voucher accessions of the same species may share an identical publication display name while retaining distinct underlying identities). Conversely, during cross-tree taxon alignment and bipartition matching, strict one-to-one identity correspondence is enforced: if two distinct tips in a tree collide onto the same normalized matching key, the import is rejected immediately with an explicit ambiguity error.

##### Unrooted branch split matching across inference methods

In the Phylogram workspace, a primary tree can be paired with up to 20 auxiliary support trees inferred by different methods (for example, pairing a concatenated maximum-likelihood tree with an ASTRAL coalescent species tree, Zhang et al., 2018, or a Bayesian consensus tree). Because different inference programs often output unrooted trees or root trees on different outgroup edges, matching internal nodes by rooted descendant sets can fail to pair identical branches. PhyloRender instead evaluates each tree over its normalized tip set $\mathcal{T}$ (aligning unique `out_` or `outgroup_` prefix variants automatically across trees) and indexes internal branches by their canonical unrooted taxon splits (bipartitions). Removing any internal branch $e$ partitions the tree's $N$ tips into two complementary non-empty subsets, $A_e$ and $B_e = \mathcal{T} \setminus A_e$, where trivial terminal splits with $|A_e| < 2$ or $|B_e| < 2$ are ignored. PhyloRender represents branch $e$ by the smaller subset (or by the lexicographically earlier sorted subset when $|A_e| = |B_e|$). For trees inferred over the same taxon set $\mathcal{T}$, this canonical split key is invariant to root placement, so identical evolutionary branches pair automatically regardless of how each input tree was rooted.

##### Support-scale verification and full-support classification

Branch support values are validated independently for each slash-delimited column of each source tree before rendering. In automatic scale-detection mode, a column is classified as percentage supports ($[0, 100]$) only when it contains values $> 1$ and contains no fractional values strictly within $(0, 1)$. Whenever a column contains any fractional value in $(0, 1)$ (even if mixed with values $> 1$) or contains only `0` and `1` without values $> 1$, PhyloRender rejects the ambiguous automatic scale and requires an explicit unit declaration (`probability` or `percent`). Low-support masking (replacing values strictly below a chosen cutoff with an en-dash –) and gene-tree discordance thresholds are always evaluated on raw unrounded numbers. Visual full-support styling (such as drawing subtending branches at $2.5\times$ baseline stroke width in place of repetitive `100/1.00` text) is applied only when every configured support source is present on that branch and reaches its respective maximum (`100` for percentages, `1` for probabilities) at the active display rounding precision (without rounding in raw-string `original` mode).

#### S1.2 | Score-Guided Node-Label Placement and Bridged Leader-Line Routing

##### Internal design coordinate space, traversal order, and local obstacle envelope

All geometric layout and collision-avoidance calculations execute on a fixed-width internal design canvas of $1{,}511$ px, which is linearly scaled upon completion by $1{,}000/1{,}511 \approx 0.6618$ onto PhyloRender's canonical $1{,}000$-unit (u) figure width (all pixel constants in Sections S1.2 and S1.3 refer to this $1{,}511$-px design space). In the Phylogram workspace, labels containing narrow masked markers (–) are placed before purely numeric strings (preserving tree traversal order within each group) so compact threshold markers can occupy tight slots beside their nodes before wider multi-method strings (such as `98/0.99`) are routed; in the Time Tree and Tanglegram workspaces, candidate nodes are ordered from top to bottom by vertical canvas coordinate. Around each internal node at $(x_0, y_0)$, the solver collects all branch segments, internal node markers, rendered tip-label boxes, and previously placed node labels and leader lines within a local search box extending $R = 192$ px (96 px in Tanglegram) plus the preferred label dimensions and an 8-px margin around $(x_0, y_0)$.

##### Multi-font candidate generation across geometric corridors

Rather than fixing a single offset or shrinking text arbitrarily, the engine evaluates discrete readable font sizes $s \in \{s_{\text{pref}}, s_{\text{pref}} - 1, \ldots, s_{\text{min}}\}$ stepping down in 1-px increments from the preferred node-label size to the configured readability floor. At each candidate size $s$, the exact SVG glyph bounding box $(w, h)$ of the label string is measured on the live canvas, and three families of candidate rectangles are generated around $(x_0, y_0)$. The first family samples directional radial rings across six primary orientations (left, lower-left, upper-left, right, upper-right, and lower-right) at adjacent clearance (4 px), expanding radial distance bands from 8 to 192 px (supplemented by cardinal vertical offsets and an orthogonal offset grid) with priority given to the ancestral side of the split. The second family samples branch-corridor slots by pairing horizontal coordinates aligned with the clearance edges (±3.05 px) of nearby vertical branch segments and the midpoints of corridors between parallel vertical branches against the top, midpoint, bottom, and node-clamped offsets of open vertical intervals between adjacent horizontal branches (with a 3.25-px vertical guard). The third family samples obstacle-boundary slots immediately above or below already placed neighboring labels (with a 2.05-px vertical guard) across horizontal offsets from the node, followed by spatial sector sampling across distance and quadrant bands.

##### Exact Liang–Barsky collision filtering

Every candidate rectangle is tested against all local obstacles before routing evaluation. Intersections between padded candidate text boxes (3-px clearance around branch segments, 2-px around other text boxes, 1.5-px around existing leader lines, $\max(3, r_{\text{node}}) + 1$ px around foreign internal nodes, and 4-px canvas margin clearance) and horizontal, vertical, or diagonal line segments are computed exactly using the parametric Liang–Barsky line-clipping algorithm rather than coarse bounding-box approximations. Any candidate colliding with a branch, tip label, placed node label, leader line, or foreign node is rejected immediately.

##### Joint typography and leader-routing cost function and multi-pass global refinement

For each collision-free candidate rectangle $\mathbf{r}$ at font size $s$, PhyloRender evaluates straight dashed leader-line connections from $(x_0, y_0)$ to central attachment points along the nearest visible edge of the label box. Font size and leader geometry compete simultaneously under a single additive objective function:

$$C(s, \mathbf{r}) = C_{\text{font}}(s) + C_{\text{route}}(\mathbf{r}) + C_{\text{pos}}(\mathbf{r})$$

The typography penalty $C_{\text{font}}(s)$ increases quadratically as the font size steps down from $s_{\text{pref}}$ toward $s_{\min}$:

$$C_{\text{font}}(s) = 20\,u_s + 25\,u_s^2, \quad u_s = \frac{\max(0,\ 1 - s/s_{\text{pref}})}{0.05}$$

When node-arrow leader lines are enabled, $C_{\text{pos}}(\mathbf{r}) = 0$ in the Phylogram and Time Tree workspaces (in Tanglegram, inward-facing placements add a side penalty of 18; when leader lines are disabled, $C_{\text{pos}}$ penalizes non-adjacent drift), and any leader path that intersects another text box is disallowed ($C_{\text{route}} = \infty$). For legal leader paths, the routing cost combines topological crossings, length, and angular deviation:

$$C_{\text{route}}(\mathbf{r}) = 2800\,n_{\text{node}} + 650\,n_{\text{leader}} + 180\,n_{\text{branch}} + 10\,u_L + 10\,u_L^2 + 300\max(0, u_L - 2)^2 + 6\left(\frac{|\theta - 45^\circ|}{15^\circ}\right)^2 + 100\,E(\theta, L)$$

where $L$ is the leader length in design pixels, $u_L = L/L_{\min}$ with minimum target leader length $L_{\min} = 14$ px, $\theta \in [0^\circ, 90^\circ]$ is the leader angle relative to the horizontal axis (targeting an ideal $45^\circ$ diagonal within a preferred $20^\circ$–$80^\circ$ window), $E(\theta, L) = \max(0, 19.75^\circ - \theta) + \max(0, \theta - 80.25^\circ) + \max(0, 13.9 - L)$ penalizes out-of-window angles and sub-minimum pointer lengths (within tolerances of $0.25^\circ$ and $0.1$ px), and $n_{\text{branch}}$, $n_{\text{leader}}$, and $n_{\text{node}}$ count crossings over branch lines, existing leader lines, and foreign node clearances, respectively. The solver selects $(s^*, \mathbf{r}^*)$ minimizing $C(s, \mathbf{r})$ and breaks ties within $10^{-6}$ in favor of larger font sizes. Following initial placement and live `getBBox` verification, Phylogram and Time Tree layouts run multi-pass global ink-box refinement passes (iterative collision repair, single-label and pairwise discrete slot swaps, font-size recovery, and crossed-leader quadrant uncrossing). Whenever a leader line crosses a branch segment ( $n_{\text{branch}} > 0$), a white halo underlay (3.8 px stroke width at 0.97 opacity) is rendered beneath the 1.15-px dashed pointer above the branch layer to create a clean white visual gap (“bridge”) at every branch crossing.

#### S1.3 | Hierarchical Clade Brackets, ICS Geologic Timescale Adaptation, Aspect Balancing, and Two-Panel Splitting

##### Multi-tier clade hierarchy and collinear vertical alignment

Taxonomic or phylogenetic highlights are defined semantically by one or more raw tip identifiers (a single tip highlights an individual terminal lineage, whereas two or more tips define the clade subtended by their MRCA). PhyloRender sorts defined groups by ascending tip count and derives their nesting hierarchy automatically by strict subset inclusion of descendant tip sets: an inner clade (tier $t = 0$) colors its descendant branches and tip names, while enclosing outer clades occupy outer tiers ($t = 1, 2, \ldots$). In automatic line-style mode, tier $t = 0$ uses solid brackets, tier $t = 1$ uses cap-locked evenly spaced dashed brackets, and higher outer tiers ($t \geq 2$) use dash-dot brackets with solid horizontal end-caps. Explicit tier indices ($t \in \{0, \ldots, 5\}$) are also supported so a monotypic subfamily and its sole tribe can occupy distinct nested tiers on the exact same node. Clade title font sizes default to 3 u larger than the active tip-label font size. Whenever two or more tiers are present (or whenever tips are flush-aligned, as in chronograms and cloudograms), all brackets in tier $t$ are locked onto a shared horizontal coordinate $X_t$ to form strictly aligned vertical columns; if two vertical titles in the same tier come within 8 px vertically, only the colliding text is shifted rightward into an adjacent lane in steps of $\mathrm{round}(1.35\, s_{\mathrm{clade}})$ while the bracket bars remain collinear on $X_t$.

##### Official ICS geologic timescale and adaptive narrow-cell fitting

The Time Tree and Cloudogram workspaces share a single geologic background and axis renderer backed by the official International Commission on Stratigraphy (ICS `2026/06`) chronostratigraphic chart spanning 0–4567 Ma (Cohen et al., 2013), including formal Precambrian eras and periods. For each visible interval $[t_{\mathrm{young}}, t_{\mathrm{old}}]$ of pixel width $W_{\mathrm{cell}}$, the label fitter tests in order whether the full interval name (e.g., `Pleistocene`), standard short abbreviation (`Pst.`), or shortest abbreviation (`Ps`) fits at the default period or epoch font size $s_0$ with internal horizontal padding. When a narrow interval near the present (such as Quaternary `Q`, Pliocene `Pl`, or Pleistocene `Ps` on a 100 Ma axis) cannot accommodate even its shortest abbreviation at $s_0$, PhyloRender scales only that cell's shortest abbreviation down to $s_{\mathrm{fit}} = \min(s_0, s_0 \cdot W_{\mathrm{avail}}/W_{\mathrm{natural}})$ (where $W_{\mathrm{avail}} = W_{\mathrm{cell}} - \min(4, \max(2, 0.18\, W_{\mathrm{cell}}))$, stepping down in 0.25-px increments until the measured glyph width fits inside the cell). Microscopic intervals whose available width is non-positive or whose fitted abbreviation would fall below the 4-px readability floor retain their colored stratigraphic band and hover metadata but omit printed text so labels never spill across borders or require external arrows.

##### Median-based tip-width compression and portrait row expansion

To prevent a few unusually long taxon or voucher strings from compressing the horizontal tree span, PhyloRender measures the natural glyph widths $\{w_1, \ldots, w_N\}$ of all displayed tip labels and computes their median $W_{\mathrm{med}}$. Only outlier labels exceeding $1.25\, W_{\mathrm{med}}$ are compressed horizontally (via SVG `textLength` with `spacingAndGlyphs`) to cap their width at $1.25\, W_{\mathrm{med}}$; shorter names remain at natural proportions without character truncation or changes in font height. Vertically, in the Phylogram, Time Tree, and Cloudogram workspaces, automatic portrait balancing expands row spacing on sparse trees ($N \leq 80$) toward a target height-to-width aspect ratio of 1.25, bounded above by min(3 em, 72 px) ($\approx 47.65$ u). Across $80 < N < 120$ tips, the extra row expansion blends smoothly back to the compact baseline spacing via the cubic Hermite weight $w(p) = 1 - 3p^2 + 2p^3$, where $p = \mathrm{clamp}((N - 80)/40, 0, 1)$.

##### Automated monophyletic two-panel splitting

For tall phylograms and chronograms exceeding the single-panel capacity ($N > 76$ tips), two-panel export splits the tree once after outgroup rooting and ladderization. All tips in the rooted outgroup are marked as protected so the root and outgroup remain in Panel 1. Among non-root internal nodes containing no outgroup tips, the splitter identifies candidate monophyletic clades of size $K$ that leave at least $N_{\mathrm{min}} = \max(2, \min(38, \lceil 0.25N \rceil))$ overview tips in Panel 1. Candidates satisfying $K \geq N - 75$ (which bring Panel 1, including one continuation guide row, within the 76-row capacity) are ranked by bottom-most terminal index, breaking ties among nested clades sharing the bottom tip in favor of the smallest sufficient clade $K$ so Panel 1 retains maximal overview context. Panel 1 replaces the extracted subtree with a continuation guide row indicating the continued $K$-tip clade in Panel 2 (preserving the stem branch length on phylograms, or extending to the present 0 Ma line on chronograms), while Panel 2 renders the complete $K$-tip monophyletic clade at the identical 1,000-u canvas width and font scale and inherits any spanning outer clade bracket automatically.

#### S1.4 | Tanglegram Sister-Clade Untangling Algorithm

##### One-to-one taxon audit and degree-2 branch-length preservation

In the Tanglegram workspace, two trees $T_1$ (left, primary reference) and $T_2$ (right, comparison) are matched one to one by normalized raw tip identities (trimmed of outer quotes, collapsed across whitespace/underscore runs, and case-folded, rejecting intra-tree duplicate collisions). If the two tip sets $\mathcal{L}(T_1)$ and $\mathcal{L}(T_2)$ are not identical, equal-count name mismatches prompt source-name verification first, while differing taxon sets require an explicit retention policy: `intersection` prunes both trees to $\mathcal{L}(T_1) \cap \mathcal{L}(T_2)$ while summing serial branch lengths across collapsed non-root degree-2 internal nodes so pairwise patristic distances remain exact, whereas `union` retains all tips from both trees and draws connectors exclusively across shared taxa.

##### Single-pass median-rank sister-clade untangling on the comparison tree

To preserve the user's chosen reference ordering, PhyloRender keeps the vertical clade order of the left primary tree $T_1$ fixed (as determined by its ladderization setting or manual sister-clade swaps) and rotates sister clades on the right comparison tree $T_2$ only. Each retained tip $t$ in $T_1$ occupies a fixed vertical row index $r_1(t) \in \{0, 1, \ldots, |\mathcal{L}(T_1)|-1\}$. Traversing $T_2$ bottom-up in postorder, each child subtree $c_i$ of an internal node $u$ is assigned the median row index in $T_1$ of its shared descendant tips:

$$m(c_i) = \text{median}\{\, r_1(t) \mid t \in \mathcal{L}(c_i) \cap \mathcal{L}_{\text{shared}} \,\}$$

(with $m(c_i) = +\infty$ under `union` if $c_i$ contains no shared tips, placing unmatched subtrees after shared siblings while preserving their relative input order). Sorting the children of every internal node $u$ in $T_2$ by ascending $m(c_i)$ (preserving original order on ties) yields a candidate topology-preserving orientation of $T_2$, which is accepted if and only if it does not increase the shared-taxon connector inversion count relative to the unmodified orientation:

$$K = \sum_{1 \le i < j \le M} \mathbb{1}(r_2(\pi_i) > r_2(\pi_j))$$

where $(\pi_1, \ldots, \pi_M)$ is the top-to-bottom sequence of shared taxa in $T_1$ and $r_2(\pi_i)$ is the vertical rank of $\pi_i$ in $T_2$. Shared tips are joined across the central corridor of net width $W = \text{right} - \text{left}$ between the two tip-label columns by horizontal-entry cubic Bézier curves (`M x1 y1 L left y1 C left+0.36W y1, right−0.36W y2, right y2 L x2 y2`) so connectors never cross label text.

#### S1.5 | Built-in Gene-Tree Discordance, Penalized-Likelihood Visual Alignment, and Cloudogram Rendering

##### Gene-tree outgroup rooting and order-invariant four-category discordance counting

In the Cloudogram workspace, a rooted species tree $T_S$ (taxon set $\mathcal{S}$, up to 1,000 taxa) is analyzed alongside a batch of gene trees $\{G_1, \ldots, G_M\}$ (up to 20,000 trees per run). Under the default outgroup-rooting policy, using the tip set $\mathcal{O}_{\text{root}}$ of one root child of $T_S$, each gene tree $G_k$ is rooted automatically on the separating edge of its observed outgroup subset $\mathcal{O}_{\text{root}} \cap \mathcal{L}(G_k)$ whenever that subset is non-empty, leaves at least two ingroup taxa, and forms a single monophyletic split in the unrooted gene tree; non-conforming gene trees are logged individually in the audit report and excluded without discarding the remaining collection.

Following Smith et al. (2015), every internal node $v$ of $T_S$ defines a rooted ingroup clade $I_v \subset \mathcal{S}$ and an outside taxon set $O_v = \mathcal{S} \setminus I_v$. For each comparable rooted gene tree $G_k$ with observed taxon set $S_k \subseteq \mathcal{S}$ (encoded as arbitrary-precision `BigInt` bitmasks), let $I_{v,k} = I_v \cap S_k$ and $O_{v,k} = O_v \cap S_k$ denote the projection of node $v$ onto $S_k$. Given a support cutoff $\tau \in [0, 0.999]$, let $\mathcal{C}(G_k, \tau)$ denote all non-trivial internal clades of $G_k$ when $\tau = 0$ (no support filter, retaining unannotated clades as well), or all internal clades carrying a numeric support strictly exceeding $\tau$ when $\tau > 0$. At species-tree node $v$, gene tree $G_k$ is **uninformative** if $|I_{v,k}| < 2$ or $|O_{v,k}| < 1$, or if no clade in $\mathcal{C}(G_k, \tau)$ either matches or conflicts with $I_{v,k}$. When missing taxa in $G_k$ cause multiple nested ancestor nodes of $T_S$ to project onto the exact same subset $I_{v,k}$, ownership of $I_{v,k}$ is assigned uniquely to the most specific (smallest-$|I_v|$) species-tree node: $G_k$ is counted as **concordant** at $v$ if $I_{v,k} \in \mathcal{C}(G_k, \tau)$ and $v$ is that unique most-specific owner (preventing missing data from inflating concordance on deeper backbone nodes, which are recorded as uninformative). Conversely, $G_k$ is counted as **conflicting** at $v$ if $\mathcal{C}(G_k, \tau)$ contains one or more clades $C \subset S_k$ topologically incompatible with $I_{v,k}$ ($C \cap I_{v,k} \neq \emptyset$, $I_{v,k} \setminus C \neq \emptyset$, and $C \cap O_{v,k} \neq \emptyset$).

When all gene trees have complete taxon sampling ($S_k = \mathcal{S}$), conflicting clades group by exact bitmask equality, separating the **dominant alternative** from **remaining alternatives**. When some gene trees lack taxa ($S_k \subset \mathcal{S}$), sequential greedy binning would depend on gene-tree input order. PhyloRender eliminates order dependence by pairing each observed conflicting clade with its observed tree scope $(C, S_k)$ and forming candidate clades $(C^*, S^*)$ where $S^* = \cup_{(C^*, S_j)} S_j$ is the union of scopes across trees directly observing $C^*$. A partial conflicting split $(C, S_k)$ contributes to $(C^*, S^*)$ if and only if $S_k \subseteq S^*$, $C = C^* \cap S_k$, and $(S_k \backslash C) \subseteq (S^* \backslash C^*)$, with each conflicting gene tree contributing at most one count to any candidate. Taking the maximum pooled count $n_{\text{dom}}(v)$ across candidates and setting $n_{\text{other}}(v) = n_{\text{conflict}}(v) - n_{\text{dom}}(v)$ guarantees strict invariance to gene-tree input order.

#### Penalized-likelihood ultrametric visual alignment

To overlay gene trees visually onto the time axis of $T_S$, PhyloRender selects comparable gene trees containing the complete species set ($\mathcal{L}(G_k) = \mathcal{S}$) and valid ML substitution branch lengths (excluding degenerate trees with $\geq 80\%$ zero-length terminal branches, bounded at $30,000$ total overlay nodes without subsampling) and scales each tree to shared species-tree clade ages (an undated ML species tree can also be calibrated to a user-supplied root age by the same solver). Following Sanderson (2002) and Smith & O'Meara (2012), each non-root branch $i \in \{1, \ldots, |E|\}$ with parent $p(i)$ and ML branch length $b_i \geq 0$ is mapped to an effective character count $c_i = \text{round}(\max(b_i, 1/N_{\text{sites}}) \cdot N_{\text{sites}})$ with $N_{\text{sites}} = 10^8$ and smoothing weight $\lambda = 100$. Terminal nodes have $t_i = 0$, the root has $t_0 = T_{\text{root}}$, and every internal gene-tree node matching a species-tree clade has fixed age $t_i = t_i^{\text{fixed}}$. The rate-smoothed Poisson penalized-likelihood objective is:

$$\mathcal{J}(\mathbf{r}, \mathbf{t}) = \frac{1}{N_{\text{sites}}}\left[\sum_{i=1}^{|E|} (r_i d_i - c_i \ln(r_i d_i)) + \lambda \sum_{p(i) \neq 0} (r_i - r_{p(i)})^2 + \frac{\lambda}{k_0} \sum_{j \in \text{children}(0)} (r_j - \bar{r}_0)^2\right]$$

where $r_i > 0$ is the branch rate, $d_i = t_{p(i)} - t_i > 0$ is the branch duration, $k_0$ is the root degree, and $\bar{r}_0 = \frac{1}{k_0} \sum_{j \in \text{children}(0)} r_j$.

To minimize $\mathcal{J}$ without external C++ `ADOL-C` or `NLopt` dependencies, PhyloRender maps the constrained variables onto an unconstrained vector $\mathbf{x} \in \mathbb{R}^{1+|E|+|V_{\text{free}}|}$. Branch rates are parameterized in log space as $r_i = \exp(\text{clamp}(x_0 + \alpha x_i, -45, 45))$, with scale factor $\alpha$ normalized from total effective substitutions and initial total branch duration. For each free internal node $u \in V_{\text{free}}$, a bottom-up pass computes its oldest fixed descendant bound $L_u = \max(\{0\} \cup \{t_w^{\text{fixed}} \mid w \in \text{desc}(u)\})$, and a top-down pass expresses $t_u$ as a logistic interpolation between its parent age $t_{p(u)}$ and $L_u$:

$$t_u = L_u + \sigma(z_u)(t_{p(u)} - L_u), \quad \sigma(z_u) = \frac{1}{1 + e^{-z_u}}, \quad z_u \in [-32, 32]$$

Keeping $0 < \sigma(z_u) < 1$ strictly in double-precision arithmetic guarantees positive branch durations $d_i > 0$ even across zero-length or micro-length radiation edges. Exact $\mathcal{O}(|V|)$ analytical gradients are obtained via postorder backpropagation:

$$\frac{\partial \mathcal{J}}{\partial z_u} = \frac{\partial \mathcal{J}}{\partial t_u}(t_{p(u)} - L_u)\sigma(z_u)(1 - \sigma(z_u)), \quad \frac{\partial \mathcal{J}}{\partial t_{p(u)}} \leftarrow \frac{\partial \mathcal{J}}{\partial t_{p(u)}} + \frac{\partial \mathcal{J}}{\partial t_u}\sigma(z_u)$$

and minimized by a built-in deterministic L-BFGS solver (history size $m = 16$, two-loop recursion with Barzilai–Borwein Hessian scaling, regular Wolfe line search with $c_1 = 10^{-4}$ and $c_2 = 0.9$, and gradient infinity-norm tolerance $10^{-11}$).

#### Cloudogram row anchoring, species-tree MRCA coloring, and deterministic jitter

During cloudogram layout, every gene-tree internal node whose descendant taxon set matches a clade of $T_S$ is anchored to that species-tree node's exact vertical coordinate ($Y$); gene-specific discordance nodes initialize at the mean $Y$ of their children and, when spanning $\geq 3$ rows, undergo two local refinement sweeps across quantile offsets to minimize branch crossings and close-approach crowding before settling near the child midpoint. Each gene branch is colored in $\mathcal{O}(\log N)$ time via a binary-lifting lowest common ancestor table on $T_S$, inheriting the group color of its descendant tips' species-tree MRCA (uncolored backbone ancestors remain neutral gray). When optional branch jitter (strength $\in \{1, \ldots, 100\}$, default 0) is enabled, non-zero-span gene branches are drawn as cubic Bézier curves with control points $(x_p + \Delta x/3,\ y_p + \Delta y/3 + \delta_y)$ and $(x_p + 2\Delta x/3,\ y_p + 2\Delta y/3 + \delta_y)$, where $\delta_y$ is derived via a deterministic 32-bit FNV-1a hash of the root, parent, and child topological clade signatures and tree index such that the mid-curve peak deflection reaches up to 0.5 row heights at strength 100 (corresponding to $|\delta_y| \leq \frac{2}{3}$ row heights, clamped to the plot bounds) while keeping all node endpoints and time coordinates fixed.

#### S1.6 | Shared Headless Runtime, Optimistic Concurrency Control, and Transactional State Consistency

##### Single-engine equivalence across browser and programmatic execution

To guarantee identical geometry, font metrics, and data validation across interactive and automated sessions, PhyloRender avoids maintaining separate plotting implementations for the web interface and the Model Context Protocol (MCP; Anthropic, 2024) server. Instead, the MCP server hosts the exact same modular JavaScript parsing, scientific preparation, collaborative render-scheduling, and SVG/PNG serialization code inside a managed headless Chromium runtime on the shared `1,000-u` figure contract. Batch gene-tree parsing, discordance counting, and penalized-likelihood alignment execute inside dedicated Web Workers that are terminated immediately upon cancellation, timeout, or page exit, while bounded server job queues and process-group supervision govern auxiliary local services and optional native helpers so aborted tasks leave no orphaned background processes.

##### Immutable revision history, optimistic concurrency, and atomic rollback

Every figure managed by the server maintains an append-only chain of immutable revisions (`revision 1, 2, ...`), storing its validated specification, SHA-256 hashes of all raw tree and table inputs, layout collision diagnostics, and topology fingerprints. Programmatic modification calls must supply the caller's known `base_revision` inside a per-figure serial queue; stale `base_revision` values are rejected immediately with a structured `REVISION_CONFLICT` error rather than overwriting concurrent updates. Across both interfaces, full specification imports and multi-operation edits execute atomically: the browser runtime captures a complete document, control, and cache snapshot prior to execution and restores it cleanly if any validation or rendering step fails, while the MCP store stages renders inside temporary directories and commits new revisions via atomic directory rename only after all checks succeed.

### Supplementary Methods 2: User Manual, Configurable Options Across Workspaces, and Operational Workflows

**Article Title:** *PhyloRender: an interactive and agent-ready package for reproducible phylogenetic visualization*
**Journal:** *Molecular Ecology Resources* (Software Note)
**Authors:** Jian He, Jingjing Chen, Zihao He, Jiamin Xiao, and Lei Xie

This user guide explains how to prepare publication-ready phylogenetic figures in PhyloRender, what options can be adjusted in each of the four workspaces, and how every setting maps across the interactive **Web Workbench**, the **Model Context Protocol (MCP; Anthropic, 2024) server**, and natural-language requests guided by the **Agent Skill** (`phylorender-mcp`). Underlying mathematical formulations and runtime architecture are documented separately in Supplementary Methods 1.

#### S2.1 | Quick-Start Overview: Choosing a Workspace, Interface, and Coordinate Unit

##### Selecting the right workspace for your evolutionary question

PhyloRender provides four dedicated workspaces in the top navigation bar of the Web Workbench (corresponding to the top-level `workspace` field in MCP figure specifications). Each workspace exposes the controls valid for that figure type and blocks inapplicable controls so chronological or topological data cannot be altered by mistake (Table S2.1).

**Table S2.1.** Overview of the four PhyloRender workspaces, input requirements, rooting behavior, and intentionally blocked controls.

| Workspace (`workspace`) | Primary Use Case | Required & Optional Tree Inputs | Rooting Policy | Output Width Cap & Supported Exports | Intentionally Blocked Controls (Data Safeguards) |
|---|---|---|---|---|---|
| **Phylogram** (`"phylogeny"`) | Standard ML, Bayesian, or coalescent trees; combining supports across methods (e.g., ML bootstrap + ASTRAL LPP, Zhang et al., 2018). | 1 primary Newick/Nexus tree (`main_tree`, up to 1,500 tips); optional up to 20 support trees (`support_trees`) and 20 node-metric trees (`other_data_trees`). | User-selected outgroup (`rooting.outgroup`), auto-root on unique `Euptelea pleiosperma` if omitted, or `outgroup: null` to keep input root. | Up to `400 mm`; `SVG`, `PNG`, two-panel `Panels`, `.json` spec. | Time axes, HPD bars, cloud layers, and two-tree connectors are hidden. |
| **Time Tree** (`"time"`) | Fossil-calibrated chronograms with divergence ages, 95% HPD credible bars, and ICS geologic bands (Cohen et al., 2013). | 1 dated summary chronogram (`main_tree`, e.g., BEAST MCC, StarBEAST3, or MCMCtree output; up to 1,500 tips); optional `other_data_trees`. | Input root and ages preserved strictly; rerooting disabled. | Up to `400 mm`; `SVG`, `PNG`, two-panel `Panels`, `.json` spec. | Outgroup rerooting, branch-support labels, support dots, full-support branch thickening, and `support_trees` are blocked so ages and HPD bars stay unambiguous. |

| Workspace (workspace) | Primary Use Case | Required & Optional Tree Inputs | Rooting Policy | Output Width Cap & Supported Exports | Intentionally Blocked Controls (Data Safeguards) |
|---|---|---|---|---|---|
| **Tanglegram** ("conflict") | Side-by-side topological comparison (e.g., nuclear vs. plastid/mitochondrial trees, or ML vs. coalescent topologies). | 1 left primary tree (main_tree) + 1 right comparison tree (comparison_tree, up to 1,500 tips); optional group palette (sample_groups). | Web preserves input roots and rotates right tree; MCP also supports rooting.outgroup to sync-root both trees on a shared outgroup side. | Up to 800 mm (double-wide); SVG, PNG, .json spec. | Horizontal branch stretching, right-margin clade bracket bars, and two-panel splitting are disabled (MCP clades[] map to group colors without bars). |
| **Cloudogram & Discordance** ("cloud") | Dated species tree overlaid with multi-locus ML gene trees and 2-part or 4-part conflict pies (following Smith et al., 2015). | 1 species tree (main_tree) + 1 multi-tree gene-tree file (gene_trees, up to 20,000 trees; complete-taxon ML subset up to 30,000 nodes scaled via built-in L-BFGS aligner following Sanderson, 2002; Smith & O'Meara, 2012). | Species-tree root preserved; gene trees rooted by species-tree outgroup (phyparts_outgroups) or kept pre-rooted ("input"). | Up to 400 mm; SVG, PNG, .json spec; node discordance TSV table. | Species-tree rerooting, numeric support strings, and two-panel splitting are disabled (horizontal stretching is hidden in Web UI, available in MCP). |

#### How the Web Workbench, MCP Server, and Agent Skill work together

In the **Web Workbench**, controls are organized into up to seven left-hand settings tabs (**Data**, **Layout**, **Tip Labels**, **Annotations**, **Time Scale** in *Time Tree*/*Cloudogram*, **Clades** outside *Tanglegram*, and **Export**), where all bounded numeric settings use compact − / number / + steppers without sliders, paired with a bottom **Optimize Node Label Placement** button and direct canvas interactions (clicking branches to define clades, double-clicking tip names to rename display labels, dragging node labels, right-clicking branches to reroot in *Phylogram*, or right-clicking left-tree nodes to swap sister clades in *Tanglegram*). In the **MCP server**, AI assistants create and revise the exact same figure state through 15 structured tools (create_figure, edit_figure, export_figure, etc.). When the companion **Agent Skill** is active in an MCP client, users state requests in plain language (for example, *"Root on Dipteronia, merge ASTRAL supports onto the ML tree, and add solid tribe brackets inside dashed subfamily brackets"*), and the assistant translates them into validated MCP calls.

#### Understanding PhyloRender's coordinate and print units

Every PhyloRender figure is laid out on a canonical canvas width of **1,000 drawing units (u)**, projected linearly from a 1,511-px internal reference canvas (1 px ≈ 0.6618 u). Canonical typography, stroke, and marker dimensions use *_u fields (for legacy compatibility, *_pt fields convert at **1 pt = 4000 / 4533 u ≈ 0.8824 u**, and legacy pixel fields convert at 1 px ≈ 0.6618 u). Physical print width (options.output.width_mm, where 0 selects the workspace maximum of 400 mm or 800 mm) and raster density (options.output.dpi, 72–1200 DPI, default 300 DPI in both Web and MCP) scale only the exported physical width or pixel count. Increasing DPI sharpens PNG exports for journal submission without shrinking fonts or shifting label placements.

### S2.2 | Shared Configurable Options Across Workspaces

The four tables below list all general layout, typography, clade-grouping, and export controls shared across workspaces, showing each control's default and valid range, its Web Workbench location, its exact MCP specification path and edit_figure operation (op), and an example natural-language Agent Skill prompt.

##### S2.2.1 | Tree layout, spacing, and stroke weight

**Table S2.2a.** Shared tree geometry, spacing, ladderization, and scale-bar controls across the three interfaces.

| Adjustable Feature | Default & Valid Range | Web Workbench Location | MCP Spec Field / `edit_figure` Operation | Example Natural-Language Prompt (Agent Skill) |
|---|---|---|---|---|
| **Sister-clade ordering (ladderization)** | `small-top` (`file`, `small-top`, `large-top`) | `Layout` → *Sister-Clade Ordering* (*Primary Ordering & Tangle Minimization* in *Tanglegram*) | `options.layout.ladderize` / `op: "set_options"` | *"Put larger sister clades on top"* or *"Keep the original sister-clade order from the file."* |
| **Auto-balanced portrait tip spacing** | `true` (targets `H/W ≈ 1.25` up to `min(3 em, 47.65 u)` on ≤ 80 tips, fading to compact at ≥ 120 tips; not applied to *Tanglegram*) | `Layout` → *Tree Span & Spacing* (`Auto-balance tip spacing & portrait aspect ratio`) | `options.layout.automatic_tip_spacing` (`true` / `false`) / `op: "set_options"` | *"Keep default automatic tip spacing"* or *"Turn off automatic portrait row expansion."* |
| **Manual vertical tip spacing** | Auto by glyph height (`2.98–47.65 u` / legacy `4.5–72 px`, or `0.9–4.0 em` relative to tip font size) | `Layout` → *Tree Span & Spacing* (`Vertical tip spacing · u` stepper) | `options.layout.tip_spacing_u` or `tip_spacing_em` / `op: "set_options"` | *"Increase vertical tip spacing to 1.6 em so rows have more breathing room."* |
| **Horizontal branch span** | `100%` (`25–400%`; visible in *Phylogram* and *Time Tree* Web UI; also supported in *Cloudogram* via MCP) | `Layout` → *Tree Span & Spacing* (`Horizontal branch span · %` stepper) | `options.layout.horizontal_scale` / `op: "set_options"` | *"Stretch the horizontal branch span to 140% to open up short internal branches."* |
| **Branch stroke width** | `0.79 u` (`0.33–2.65 u`, corresponding to `0.5–4.0` design `px`; visible in *Phylogram*/*Time Tree* Web UI, all modes in MCP) | `Layout` → *Branch Scaling & Stroke* (`Branch stroke width · u` stepper) | `options.layout.branch_width_u` / `op: "set_options"` | *"Set branch line thickness to 1.0 drawing units."* |
| **Scale-bar typography** | Arial, `11.91 u` (`4.63–15.88 u`), `#26302b`, upright (*Phylogram* substitution scale bar) | `Layout` → *Scale Bar Typography* (`Font family`, `Text color`, `Font size · u`, `Bold`, `Italic`) | `options.typography.scale` (`family`, `size_u`, `color`, `bold`, `italic`) / `op: "set_options"` | *"Make the substitution scale bar text 10.5 units and bold."* |

##### S2.2.2 | Tip labels, taxonomic nomenclature, and display renaming

**Table S2.2b.** Terminal taxon typography, binomial italicization, and non-destructive display renaming across the three interfaces.

| Adjustable Feature | Default & Valid Range | Web Workbench Location | MCP Spec Field / `edit_figure` Operation | Example Natural-Language Prompt (Agent Skill) |
|---|---|---|---|---|
| **Tip font family** | `Arial` (`Arial`, `Libre Baskerville`, `Source Serif 4`, `Arimo`, `Noto Sans SC`) | `Tip Labels` → *Tip Font & Color* (`Font family`) | `options.typography.tips.family` / `op: "set_options"` | *"Switch tip labels to Source Serif 4 (Times style)."* |

| Adjustable Feature | Default & Valid Range | Web Workbench Location | MCP Spec Field / `edit_figure` Operation | Example Natural-Language Prompt (Agent Skill) |
|---|---|---|---|---|
| **Tip font size & bold weight** | `13.24 u` in *Phylogram*/*Tanglegram*; `15.88 u` in *Time Tree*/*Cloudogram* (`5.29–19.85 u`); `bold: false` | `Tip Labels` → *Tip Font & Color* (`Font size · u` stepper, `Bold` checkbox) | `options.typography.tips.size_u`, `bold` / `op: "set_options"` | *"Set tip font size to 14 units and make all tip names bold."* |
| **Tip color & clade color inheritance** | `#17211b` (`#202a30` in *Cloudogram*); `follow_clade_color: true` | `Tip Labels` → *Tip Font & Color* (`Text color`, `Match assigned clade color`) | `options.typography.tips.color`, `follow_clade_color` / `op: "set_options"` | *"Keep tip names black instead of inheriting clade colors."* |
| **Scientific binomial italicization** | `auto` (`auto` italicizes Latin genus/epithets and keeps vouchers upright; `all`; `none`) | `Tip Labels` → *Taxonomic Nomenclature* (`Italicization rule`) | `options.typography.tips.italic_mode` (`"auto"`, `"all"`, `"none"`) / `op: "set_options"` | *"Auto-italicize Latin species names while keeping voucher numbers upright."* |
| **Infraspecific rank standardization** | `true` (normalizes `SSP`/`subspecies` → `subsp.`, `VAR`/`variety` → `var.`, `forma` → `f.` in roman type) | `Tip Labels` → *Taxonomic Nomenclature* (`Standardize infraspecific rank abbreviations`) | `options.typography.tips.normalize_ranks` (`true` / `false`) / `op: "set_options"` | *"Standardize subspecies and variety rank abbreviations."* |
| **Custom tip display names (single or batch mapping)** | Raw tree names shown by default; overrides (up to 5,000) change display text only without altering matching IDs | Double-click any tip on canvas, or `Tip Labels` → *Rename Tip Labels* (`Upload Mapping Table` CSV/TSV/TXT/`.xlsx`) | `tip_label_overrides` (`{raw_tip: display_name}`) / `op: "set_tip_label"`, `"remove_tip_label"`, `"merge_tip_labels"`, `"replace_tip_labels"`, `"clear_tip_labels"` | *"Apply specimen display names from `voucher_map.csv` without changing underlying taxon IDs."* |
| **Long tip-label width cap** | Automatic cap at `1.25×` median natural tip width (compresses outliers horizontally without truncation) | Automatic across all four workspaces | Enforced automatically by shared figure contract | Applied automatically; no manual tuning needed. |

#### S2.2.3 | Clade brackets, multi-tier hierarchy, and lineage coloring

**Table S2.2c.** Semantic MRCA clade grouping, nested bracket tiers, rich-text clade segments, and bracket typography (bracket bars rendered in *Phylogram*, *Time Tree*, and *Cloudogram*; see Section S2.3.3 for *Tanglegram* group palettes).

| Adjustable Feature | Default & Valid Range | Web Workbench Location | MCP Spec Field / `edit_figure` Operation | Example Natural-Language Prompt (Agent Skill) |
|---|---|---|---|---|
| **Add, update, or remove a clade group** | Up to 200 groups defined by `mrca_of` ( ≥ 2 raw tips lock an MRCA clade; 1 tip highlights a single lineage) | `Clades` → *Clade & Taxon Highlighting* (`Select Clade or Taxon`, click branch/tip; `Clear All` clears all) | `clades[]` (`mrca_of`, `name`, `color`, `clade_id?`) / `op: "add_clade"`, `"update_clade"`, `"remove_clade"` (or `update_figure` with `clades: []`) | *"Highlight Sapindoideae as the MRCA of Koelreuteria_paniculata and Litchi_chinensis in teal `#2b6f6b`."* |
| **Nested bracket tier (e.g., tribe inside subfamily)** | Omitted/`null` derives tier automatically from subset nesting; explicit integer `0–5` (`0` = innermost tier) | Clade dialog → `Bracket tier` (`Automatic` or `Inner tier 1` to `Outer tier 6`) | `clades[].tier` (`0–5` or `null`/omitted) / `op: "add_clade"`, `"update_clade"` | *"Put tribes on inner tier 0 and subfamilies on outer tier 1, including monotypic Xanthoceroideae."* |

| Adjustable Feature | Default & Valid Range | Web Workbench Location | MCP Spec Field / `edit_figure` Operation | Example Natural-Language Prompt (Agent Skill) |
|---|---|---|---|---|
| **Bracket line style** | `auto` (`solid` on tier `0`, `dashed` on tier `1`, `dash-dot` on tiers ≥ 2), or explicit `solid`, `dashed`, `dash-dot` | Clade dialog → `Bracket line style` | `clades[].bar_style` (`"auto"`, `"solid"`, `"dashed"`, `"dash-dot"`) | *"Use solid lines for tribes and dashed lines for subfamilies."* |
| **Clade label orientation** | `auto` (horizontal on 1-tier phylogram if room permits; vertical on aligned or ≥ 2-tier brackets), `horizontal`, or `vertical` | Clade dialog → `Label orientation` | `clades[].label_orientation` (`"auto"`, `"horizontal"`, `"vertical"`) | *"Orient all subfamily and tribe bracket names vertically."* |
| **Bracket column alignment** | `auto` (tracks tip lengths on 1-tier phylograms; flush shared column when ≥ 2 tiers or tip-aligned), `nearby`, or `aligned` | `Clades` → *Clade Bar Alignment* (`Bar alignment mode`) | `options.layout.clade_bar_alignment` (`"auto"`, `"nearby"`, `"aligned"`) / `op: "set_options"` | *"Align all clade brackets flush along a shared right column."* |
| **Clade title font & relative size** | Arial, bold, `follow_tip_size: true` (3 u larger than active tip font size; manual `5.29–23.16 u` fixes size) | `Clades` → *Clade Label Typography* (`Automatic: 3 u larger than tip labels` or `Font size · u` stepper) | `options.typography.clades` (`family`, `size_u`, `follow_tip_size`, `bold`, `italic`) / `op: "set_options"` | *"Keep clade titles 3 units larger than tip names"* or *"Fix clade title font size at 18 units."* |
| **Mixed roman/italic segments within one clade label** | Uniform style by default; up to 60 styled runs ( ≤ 120 total chars, segment `size_u` `5.29–19.85 u`) | Clade dialog → *Rich-text segment styling (optional)* (`Split by words` / `+ Add segment`) | `clades[].name_segments[]` (`text`, `italic`, `bold`, `family`, `size_u`, `color`) | *"Label the clade with italic 'Clematis' followed by upright 'sect.' and italic 'Atragene'."* |

#### S2.2.4 | Export formats, print dimensions, two-panel splitting, and global re-optimization

**Table S2.2d.** Vector/raster exports, physical dimensions, two-panel pagination, portable specifications, and global label re-optimization.

| Adjustable Feature | Default & Valid Range | Web Workbench Location | MCP Spec Field / Tool Call | Example Natural-Language Prompt (Agent Skill) |
|---|---|---|---|---|
| **Vector SVG & high-DPI PNG export** | `SVG` (bundled fonts embedded as subsets; system Arial referenced directly); `PNG` (`72–1200 DPI`, default `300 DPI`) | `Export` → *Export Formats* (`Export SVG`, `Export PNG`) & *PNG Raster Resolution* (`DPI` stepper) | Set `options.output.dpi` via `op: "set_options"`, then call `export_figure` (`formats: ["svg", "png"]`) | *"Set output resolution to 600 DPI and export both SVG and PNG."* |
| **Monophyletic two-panel split** | Splits tall *Phylogram* or *Time Tree* (>76 tips) once at a monophyletic crown clade into 2 equal-width panels | `Export` → *Export Formats* (`Export Two-Panel Split`, downloads `.zip` bundle) | `export_figure` (`formats: ["panels"]`, or `["panels", "png"]` to include panel PNGs) | *"Export this 176-tip tree as a two-panel split so it fits cleanly across two columns."* |
| **Canvas aspect ratio** | `auto` (`auto`, `content`, `1:1`, `4:3`, `3:2`, `16:9`, `a4-landscape`, `a4-portrait`, or `custom` `0.40–3.00`) | `Export` → *Canvas Aspect Ratio* (`Output aspect ratio` & `Custom width / height ratio`) | `options.output.aspect_ratio`, `custom_ratio` / `op: "set_options"` | *"Use a compact fit-to-content aspect ratio."* |
| **Physical print width (mm)** | `0` = workspace cap (`0–400 mm` single-tree, `0–800 mm` *Tanglegram*) | `Export` → *Print Dimensions* (`Print width (0 = max)` stepper) | `options.output.width_mm` / `op: "set_options"` | *"Set physical print width to 180 mm for a full-page journal figure."* |

| Adjustable Feature | Default & Valid Range | Web Workbench Location | MCP Spec Field / Tool Call | Example Natural-Language Prompt (Agent Skill) |
|---|---|---|---|---|
| **Portable JSON figure specification** | Web `.json` bundles raw trees + settings; MCP `get_figure_spec` redacts raw tree text with SHA-256 hashes | `Export` → *Shared Web & AI Agent Specification* (`Export Figure Spec (.json)` / `Load Figure Spec (.json)`) | `get_figure_spec` (redacted archival JSON); populate `content` fields for direct Web `Load Figure Spec` | *"Save a complete portable figure specification JSON so I can load it in the browser workbench."* |
| **Global node-label re-optimization & font reset** | Re-runs collision-free slot and leader-line search across internal nodes; restores workspace default fonts if requested | Persistent bottom button `Optimize Node Label Placement`; `Export` → *Reset Typography* (`Restore Default Fonts & Colors`) | Executed automatically on every MCP render; `op: "reset_label_adjustments"` clears manual offsets | *"Reset manual label offsets and re-optimize all node label placements."* |

#### S2.3 | Per-Workspace Reference Guide

##### S2.3.1 | Phylogram (`workspace: "phylogeny"`)

The **Phylogram** workspace renders rooted or unrooted maximum-likelihood, Bayesian, and coalescent trees and merges branch support or node metrics from companion analyses onto the primary topology via unrooted bipartition matching (Supplementary Methods 1, Section S1.1). Users can reroot the primary tree either by selecting a single outgroup taxon in `Layout` and clicking **Reroot by Selected Outgroup**, or by right-clicking any internal branch on the canvas to reroot on a multi-taxon outgroup clade.

**Table S2.3a.** Controls exclusive to (or primarily configured within) the **Phylogram** workspace.

| Adjustable Feature | Default & Valid Range | Web Workbench Location | MCP Spec Field / `edit_figure` Operation | Example Natural-Language Prompt (Agent Skill) |
|---|---|---|---|---|
| **Outgroup rerooting (single or multi-taxon)** | Specify ≥ 1 raw tip IDs in `rooting.outgroup`, or `null` to keep input root (auto-roots on unique `Euptelea pleiosperma` if omitted) | `Layout` → *Rooting & Outgroup* (`Outgroup taxon` + `Reroot by Selected Outgroup`), or right-click branch → `Reroot at this branch` | `rooting: { outgroup: [...] }` / `op: "set_rooting"` (`outgroup: [...]` or `null`) | *"Root the tree on the two Dipteronia species (`Dipteronia_sinensis`, `Dipteronia_dyeriana`)."* |
| **Branch-length representation** | `phylogram` (`phylogram` proportional branch lengths, `cladogram` equal-depth, `aligned` tip-aligned cladogram) | `Layout` → *Branch Scaling & Stroke* (`Branch–length representation`) | `options.layout.mode` (`"phylogram"`, `"cladogram"`, `"aligned"`) / `op: "set_options"` | *"Keep proportional ML branch lengths"* or *"Display as an equal-depth cladogram."* |
| **Attach companion support trees** | Up to 20 companion trees matched by unrooted splits; discordant splits absent from a companion tree show as – | `Data` → *Supplementary Node Data* → *Add Branch Support Trees* (`Add Support Trees` & scale selector) | `support_trees[]` (`{ content, name?, source_id?, label?, field?, scale? }`) / `op: "add_support_tree"`, `"update_support_tree"`, `"remove_support_tree"` | *"Add `astral.tre` as a second support column with probability (0–1) scale."* |

| Adjustable Feature | Default & Valid Range | Web Workbench Location | MCP Spec Field / `edit_figure` Operation | Example Natural-Language Prompt (Agent Skill) |
|---|---|---|---|---|
| **Attach other node metrics (ages, rates, scalars)** | Up to 20 companion trees (also available in *Time Tree*); displays chosen annotation field without affecting support styling | `Data` → *Supplementary Node Data* → *Divergence ages, evolutionary rates & other metrics* (`Add Metric Trees`) | `other_data_trees[]` (`{ content, name?, source_id?, label?, field?, decimals? }`) / `op: "add_other_data_tree"`, `"update_other_data_tree"`, `"remove_other_data_tree"` | *"Attach substitution rate annotations (`field:rates`) from `rates.nexus` rounded to 3 decimals."* |
| **Primary support scale declaration** | `auto` (supports `auto`, `probability`, `percent`, and slash-delimited combinations; fractional `(0,1)` ambiguity requires explicit scale) | `Data` → *Primary Tree* (`Primary tree support scale`) | `options.support.scale` / `op: "set_options"` | *"The primary tree has composite SH-aLRT percentage / UFBoot percentage supports (`percent/percent`)."* |
| **Support decimal precision** | `auto` (`0` decimals for `%`, `2` decimals for `0–1` probabilities; `original` raw string; or fixed `0–6` places; shared with *Tanglegram*) | `Annotations` → *Support Decimal Precision* (`Auto by scale`, `As in file`, or `0–6` stepper) | `options.support.decimals` (`"auto"`, `"original"`, or `0–6`) / `op: "set_options"` | *"Format percentage supports as integers and posterior probabilities to 2 decimal places."* |
| **Full-support branch representation** | `branch` (`2.5×` thickened branch on full support across all columns, red dot on non-full; or `circle`, `number`, `star`, `hide`) | `Annotations` → *Support Display Conventions* (`Full support representation`) | `options.support.full_support_marker` (`"branch"`, `"circle"`, `"number"`, `"hide"`, `"star"`) / `op: "set_options"` | *"Thicken branches that reach 100% ML bootstrap and 1.00 ASTRAL LPP, and omit their `100/1.00` text."* |
| **Low-support masking cutoff** | `50%` (integer `0–100%`, evaluated per column on raw values; `0` disables masking; masked values print as –) | `Annotations` → *Support Display Conventions* (`Low-support masking cutoff · %` stepper) | `options.support.low_support_threshold` (`0–100`) / `op: "set_options"` | *"Mask individual support values below 70% as an en-dash."* |
| **Node marker dot radius** | `3.31 u` (`1.32–7.94 u`, shared with *Time Tree* and *Tanglegram*) | `Annotations` → *Node Markers* (`Node marker radius · u` stepper) | `options.layout.node_radius_u` / `op: "set_options"` | *"Reduce internal node dot radius to 2.2 units."* |
| **Node label font, preferred size & minimum floor** | Arial, `#8c2e28`, `automatic_size: true` (`preferred_size_u` `5.96–21.18 u`, compact baseline `11.91 u`; `minimum_size_u` `5.96–11.91 u`, default `6.18 u`) | `Annotations` → *Node Label Font & Color* & *Node Label Size Bounds* | `options.typography.nodes` (`family`, `color`, `preferred_size_u`, `automatic_size`, `minimum_size_u`, `bold`, `italic`) / `op: "set_options"` | *"Color node support labels dark slate `#334155` and set minimum font size floor to 7.0 units."* |
| **Automatic slot search, leader lines & per-node nudge** | `auto_placement: true`, `node_arrows: true`; individual labels can be dragged on canvas or nudged by `branch_id` | Bottom panel `Placement options & rules` checkboxes; drag any label on canvas | `options.labels.auto_placement`, `options.labels.node_arrows` / `op: "set_options"`; `op: "set_label_adjustment"`, `"remove_label_adjustment"`, `"reset_label_adjustments"` | *"Enable leader lines for crowded short-branch labels."* |

##### S2.3.2 | Time Tree (`workspace: "time"`)

The **Time Tree** workspace renders fossil-calibrated summary chronograms (such as BEAST TreeAnnotator MCC trees, StarBEAST3 species chronograms, MCMCtree outputs, or `treePL` dated trees). In the Web Workbench, importing a chronogram uses the **Import Time Tree** panel under `Data`: **Step 1** loads the dated tree file, and **Step 2** stays open automatically so you can confirm the input file's time unit (`yr`, `kyr`, or `Mya`) and choose a single uniform node-age statistic across the entire tree. Non-ultrametric trees with tip ages exceeding numerical roundoff tolerance (heterochronous tip-dated trees or sampled ancestors) are rejected clearly rather than zeroed out silently. Branch-support values and rerooting are intentionally blocked so node bars and numbers communicate divergence time and credible intervals exclusively.

**Table S2.3b.** Controls exclusive to the **Time Tree** workspace.

| Adjustable Feature | Default & Valid Range | Web Workbench Location | MCP Spec Field / `edit_figure` Operation | Example Natural-Language Prompt (Agent Skill) |
|---|---|---|---|---|
| **Source time unit & axis display unit** | `unconfirmed` until verified (`years`, `kyr`, `mya`); `display_unit: "same"` or unit conversion (`years`, `kyr`, `mya`) | `Data` → *Import Time Tree* → *Step 2 (Confirm Time Unit & Age Source)* (`Age unit in tree file`, `Axis display unit`) | `options.time.source_unit`, `display_unit` / `op: "set_options"` | *"The BEAST tree ages are in million years (`mya`); display the axis in Mya."* |
| **Uniform node-age statistic** | `auto` (`auto`, `branch_lengths`, `height`, `height_mean`, `height_median`; applied uniformly to all nodes) | `Data` → *Import Time Tree* → *Step 2* (`Node age source`) | `options.time.age_source` / `op: "set_options"` | *"Use posterior mean node heights (`height_mean`) for node coordinates and labels."* |
| **Node age visibility & decimal places** | `show_node_ages: true`; 2 decimal places (`0–3` places) | `Annotations` → *Node Ages & HPD Credible Bars* (`Show divergence ages at internal nodes`, `Node age decimal places`) | `options.time.show_node_ages`, `age_decimals` (`0–3`) / `op: "set_options"` | *"Show node divergence ages rounded to 1 decimal place."* |
| **Node age label window (dense-chronogram filter)** | On `<160` internal nodes: labels all nodes. On ≥ 160 internal nodes: labels older `50%` by default (`age_minimum: 0` labels all nodes) | `Annotations` → *Node Age Label Filter* (`Minimum age`, `Maximum age`, `Show All Node Ages`) | `options.time.age_minimum`, `age_maximum` (`null` or ≥ 0) / `op: "set_options"` | *"Show node age numbers only for divergences between 15 and 100 Mya, while keeping all HPD bars."* |
| **95% HPD credible bars & clade scope** | `show_hpd: true`; `hpd_scope: "all"` (`"all"` nodes with HPD, or `"named_clades"` defined in `Clades` only) | `Annotations` → *Node Ages & HPD Credible Bars* (`Show 95% HPD node bars`, `Nodes to display`) | `options.time.show_hpd`, `hpd_scope` (`"all"`, `"named_clades"`) / `op: "set_options"` | *"Display 95% HPD bars only on the named subfamily and tribe crown nodes."* |
| **In-bar summary tick & HPD bar styling** | `hpd_marker: "none"` (`none`, `height_mean`, `height_median`); color `#5f72a5`, opacity `40%` (`10–100%` / `0.1–1.0`), thickness `4.63 u` (`1.32–10.59 u`) | `Annotations` → *Node Ages & HPD Credible Bars* (`In-bar summary tick`, `Bar color`, `Opacity · %`, `Bar thickness · u`) | `options.time.hpd_marker`, `hpd_color`, `hpd_opacity`, `hpd_thickness_u` / `op: "set_options"` | *"Color the 95% HPD bars slate blue `#4c6a9c` at 45% opacity and thickness 4.0 units."* |
| **ICS geologic timescale backdrop** | `standard` official ICS `0–4567 Ma` palette (requires confirmed time unit), or `none` (plain white background) | `Time Scale` → *Geologic Time Scale & Axis* (`Geologic time scale backdrop`) | `options.time.geology` (`"standard"`, `"none"`) / `op: "set_options"` | *"Show the standard ICS geologic timescale bands behind the chronogram."* |

| Adjustable Feature | Default & Valid Range | Web Workbench Location | MCP Spec Field / `edit_figure` Operation | Example Natural-Language Prompt (Agent Skill) |
|---|---|---|---|---|
| **Stratigraphic & time-axis font sizes** | Independent sizes (`2.65–21.18 u` / `3–24 pt`): periods `14.56 u`, epochs `12.57 u`, axis ticks `13.90 u`, unit title `13.24 u` | `Time Scale` → *Geologic Time Scale & Axis* (`Geologic periods / eras`, `Geologic epochs`, `Axis tick numerals`, `Time unit title` steppers in `u`) | `options.time.period_size_u`, `epoch_size_u`, `axis_size_u`, `unit_size_u` / `op: "set_options"` | *"Increase time-axis tick numerals to 15 units and geologic period names to 16 units."* |

#### S2.3.3 | Tanglegram (`workspace: "conflict"`)

The **Tanglegram** workspace places a left primary tree and a right comparison tree face to face across a central cubic Bézier connector corridor, supporting double-wide exports up to `800 mm`. Each side scales independently by its own branch lengths. If the two trees have equal tip counts with unmatched names, PhyloRender prompts source-spelling verification first; when taxon sets differ, it lists every unmatched taxon and requires an explicit choice between `intersection` and `union` retention policies.

**Table S2.3c.** Controls exclusive to the **Tanglegram** workspace.

| Adjustable Feature | Default & Valid Range | Web Workbench Location | MCP Spec Field / `edit_figure` Operation | Example Natural-Language Prompt (Agent Skill) |
|---|---|---|---|---|
| **Primary & comparison trees, titles, and swap** | Titles default to `"Primary tree"` (left) and `"Comparison tree"` (right; up to `80` characters) | `Data` → *Paired Input Trees* (`Select Primary Tree`, `Select Comparison Tree`, `Swap Primary & Comparison Trees`, title inputs) | `main_tree`, `comparison_tree` (via `create_figure` / `update_figure`); `options.conflict.primary_title`, `secondary_title` / `op: "set_options"` | *"Compare `nuclear.tre` on the left titled 'Nuclear (353 loci)' against `plastome.tre` on the right titled 'Plastome'."* |
| **Taxon retention policy on mismatched trees** | Required when tip sets differ: `intersection` (retains shared taxa, summing branch lengths across collapsed nodes) or `union` (keeps all tips) | `Data` → *Taxon Set Reconciliation* (`Taxon retention policy`: `Retain shared taxa only (intersection)` vs. `Retain all taxa from both trees (union)`) | `options.conflict.sample_policy` (`"intersection"`, `"union"`) / `op: "set_options"` | *"Prune both trees to their 29 shared accessions using the intersection policy."* |
| **Left-tree ordering, manual clade swap & untangling** | Left ladderized `small-top`; right tree untangled automatically via postorder median-rank rotation without altering left order | `Layout` → *Primary Ordering & Tangle Minimization* (`Primary tree ladderization`, `Re-optimize Connector Crossings`), or right-click left node → `Swap sister clades` | `options.layout.ladderize`, `options.layout.tip_order` (for manual sister-clade swaps) / `op: "set_options"` | *"Ladderize the nuclear tree with smaller clades on top and untangle the plastid tree connectors."* |
| **Independent left & right support scales and fields** | `auto` per tree (`all`, `field:0` first value, `field:1` second value, etc. in slash-delimited `SH-aLRT/UFBoot` strings) | `Annotations` → *Support Field Selection* (`Left Primary Tree` & `Right Comparison Tree` field and scale selectors) | `options.conflict.primary_support_field`, `secondary_support_field`, `primary_support_scale`, `secondary_support_scale` / `op: "set_options"` | *"Show UFBoot (`field:1`, the second slash-delimited field) on both trees as percentages."* |

| Adjustable Feature | Default & Valid Range | Web Workbench Location | MCP Spec Field / `edit_figure` Operation | Example Natural-Language Prompt (Agent Skill) |
|---|---|---|---|---|
| **Taxon group color palette (`sample_groups`)** | Auto-detects major primary-tree clades by default; explicit groups color left clades, right tips, and central connectors | `Data` → *Clade Grouping & Palette* (`Import Taxon Group Table` CSV/TSV `sample, group, color`; `Download Group Template CSV`) | `sample_groups[]` (`{ name, color, samples: [...] }`, up to 200 groups) / `op: "set_sample_groups"`, `"clear_sample_groups"` | *"Color lineages and connectors by tribe using `sapindaceae_tribes.csv`."* |

##### S2.3.4 | Cloudogram & Discordance (`workspace: "cloud"`)

The **Cloudogram & Discordance** workspace combines up to three synchronized visual layers on one species-tree coordinate system: (1) the species-tree backbone, (2) a penalized-likelihood time-aligned ML gene-tree cloudogram colored by species-tree MRCA clades, and (3) two-part or four-part rooted gene-tree discordance pie charts at internal nodes, with an optional ICS geologic timescale backdrop. Styling changes (pie mode, opacity, jitter, clade colors, or hiding individual pies) redraw instantly from cached scientific results without rerunning discordance counting or branch scaling.

**Table S2.3d.** Controls exclusive to the **Cloudogram & Discordance** workspace.

| Adjustable Feature | Default & Valid Range | Web Workbench Location | MCP Spec Field / `edit_figure` Operation | Example Natural-Language Prompt (Agent Skill) |
|---|---|---|---|---|
| **Species-tree calibration type & root age** | `dated` preserves input ages (`dated_age_source`, `dated_unit`); undated `ml` (MCP default if omitted) requires `root_age > 0` (`root_age_unit` defaults to `"relative"`); `coalescent`/`other` draws pies only | `Data` → *Species Tree & Gene Trees* (`Species tree input type`, `Root / crown calibration age`, `Age unit`, `Node age source`, `Source time unit`) | `options.cloud.species_method`, `genes_method`, `root_age`, `root_age_unit`, `dated_age_source`, `dated_unit` / `op: "set_options"` | *"Use the dated StarBEAST3 species tree in Mya (`height_mean`) with the 86 ML gene trees."* |
| **Gene-tree rooting outgroup & policy** | `phyparts_root_policy: "outgroup"`; singleton outgroups auto-detect when `[]`, whereas multi-taxon outgroups require all tips of one species-tree root child (`"input"` keeps pre-rooted trees) | `Data` → *Gene-Tree Discordance Analysis* → *Discordance options & node table* (`Outgroup taxa for gene-tree rooting`, `Gene-tree rooting policy`) | `options.cloud.phyparts_outgroups[]`, `phyparts_root_policy` (`"outgroup"`, `"input"`) / `op: "set_options"` | *"Root the gene trees using the species-tree outgroup clade `Acer_truncatum` and `Dipteronia_sinensis`."* |
| **Discordance support cutoff & field selection** | `0%` (`0–99.9%`, no filter at `0%`, strict > on raw support when `>0%`); `last` (default) or `first` field | `Data` → *Gene-Tree Discordance Analysis* → *Discordance options & node table* (`Minimum branch support cutoff (%)`, `Multi-field support selection`, `Support scale`) | `options.cloud.phyparts_threshold`, `phyparts_field` (`"last"`, `"first"`), `phyparts_scale` / `op: "set_options"` | *"Apply a 50% bootstrap cutoff when counting concordant and conflicting gene trees."* |

| Adjustable Feature | Default & Valid Range | Web Workbench Location | MCP Spec Field / `edit_figure` Operation | Example Natural-Language Prompt (Agent Skill) |
|---|---|---|---|---|
| **Export or import discordance node table** | Built-in engine computes all 4 categories automatically; TSV node table can be downloaded or supplied | `Data` → *Gene-Tree Discordance Analysis* → *Discordance options & node table* (`Download Discordance Node Table`, `Import Precomputed Node Table`) | Top-level `phyparts_table` (`{ content, name? }`); node counts returned in `inspect_figure` report | *"Report the four-category gene-tree conflict counts at the Xanthoceras sister node."* |
| **Gene-tree cloudogram visibility, opacity & jitter** | `show_gene: true`; `opacity: 18%` (`2–60%`, baseline auto-scaled by locus count); deterministic mid-branch `jitter: 0%` (`0–100%`) | `Layout` → *Gene-Tree Cloudogram Layer* (`Show gene-tree cloudogram`, `Gene-tree stroke opacity · %`, `Cloudogram branch jitter · %`) | `options.cloud.show_gene`, `opacity` (`2–60`), `jitter` (`0–100`) / `op: "set_options"` | *"Set gene-tree cloud opacity to 14% and branch jitter to 25% so overlapping trajectories separate."* |
| **Species-tree overlay & backbone shape** | `show_species: true`; `species_shape: "triangular"` (`"rectangular"` available when cloudogram layer is hidden) | `Layout` → *Species-Tree Overlay* (`Show species tree`, `Species-tree branch geometry`) | `options.cloud.show_species`, `species_shape` (`"triangular"`, `"rectangular"`) / `op: "set_options"` | *"Hide the gene-tree cloud layer and draw the species tree as a rectangular phylogram with pies."* |
| **Discordance pie composition (`2-part` vs. `4-part`)** | `two` (`"two"`: Concordant vs. Conflicting among informative loci; `"four"`: Concordant / Top conflict / Other conflict / Uninformative) | `Annotations` → *Discordance Node Pies* (`Show node discordance pies`, `Pie composition`) | `options.cloud.show_pies`, `pie_mode` (`"two"`, `"four"`) / `op: "set_options"` | *"Switch node pies to the four-part breakdown including dominant alternative and uninformative genes."* |
| **Pie radius, legend position & legend scale** | `pie_radius_u: 21.18 u` (`5.29–35.74 u`, scales monotonically with node age); legend at `lower-left` (`auto`, `upper-left`, `middle-left`, `lower-left`) at `150%` (`75–200%`) | `Annotations` → *Discordance Node Pies* (`Maximum pie radius · u`, `Pie legend position`, `Pie legend scale · %`) | `options.cloud.pie_radius_u`, `pie_legend_position`, `pie_legend_scale` / `op: "set_options"` | *"Place the pie legend in the lower-left corner at 135% scale and set maximum pie radius to 18.5 units."* |
| **Pie age filter & hiding individual node pies** | No age filter by default; individual pies can be hidden/restored by clicking on canvas or by MRCA tip sets ( ≥ 2 tips) without changing denominators | `Annotations` → *Discordance Pie Age Filter* (`Minimum age`, `Maximum age`), or click any pie on canvas → `Hide pie chart` | `options.cloud.pie_age_minimum`, `pie_age_maximum`, `hidden_pies` / `op: "set_options"`, `"hide_cloud_pie"`, `"show_cloud_pie"`, `"reset_hidden_cloud_pies"` | *"Hide discordance pies on terminal sister pairs younger than 5 Mya to keep the shallow tips clean."* |
| **ICS geologic timescale on cloudogram** | `show_time_axis: false` by default (`true` adds top/bottom ICS stratigraphic bands using `options.time.geology` and `*_size_u` font sizes) | `Time Scale` → *Geologic Time Scale & Axis* (`Show time axis & geologic time scale on cloudogram`) | `options.cloud.show_time_axis` (`true` / `false`) + `options.time` (`geology`, `period_size_u`, `epoch_size_u`, `axis_size_u`, `unit_size_u`) / `op: "set_options"` | *"Turn on the ICS geologic timescale axis beneath the cloudogram."* |

#### S2.4 | Reference of MCP Server Tools, Atomic Edit Operations, and Agent Skill Rules

##### Complete inventory of the 15 MCP server tools and 24 `edit_figure` operations

AI assistants connected to PhyloRender interact through the 15 structured MCP tools summarized in Table S2.4. Within `edit_figure`, callers submit an atomic array of semantic `operations[]` drawn from **24 exact `op` types**: `set_options` (updates any nested `options.*` fields), `set_rooting`, five tip-renaming operations (`set_tip_label`, `remove_tip_label`, `merge_tip_labels`, `replace_tip_labels`, `clear_tip_labels`), three clade operations (`add_clade`, `update_clade`, `remove_clade`), three support-tree operations (`add_support_tree`, `update_support_tree`, `remove_support_tree`), three metric-tree operations (`add_other_data_tree`, `update_other_data_tree`, `remove_other_data_tree`), three node-label offset operations (`set_label_adjustment`, `remove_label_adjustment`, `reset_label_adjustments`), two tanglegram group operations (`set_sample_groups`, `clear_sample_groups`), and three cloud-pie visibility operations (`hide_cloud_pie`, `show_cloud_pie`, `reset_hidden_cloud_pies`).

**Table S2.4.** Functional reference of all 15 tools exposed by the PhyloRender MCP server.

| Lifecycle Stage | Tool Name | Purpose & Exact Inputs | Returned Output |
|---|---|---|---|
| **1. Pre-render tree & taxon audit** | `inspect_tree` | Parses tree text (`{ main_tree: { content, name? } }`) prior to rendering (Agent Skill clients hydrate local files into `content` locally). | Raw tip labels, branch-length profile, support-value ranges, ultrametric/HPD metadata, dense-chronogram flag ( ≥ 160 internal nodes), and topology fingerprint. |
| | `compare_tree_samples` | Compares raw tip sets (`{ main_tree, comparison_tree }`) prior to building a tanglegram. | Shared taxon count, exact `primary_only` and `comparison_only` taxon lists, equal-count mismatch flag, and allowed sample policies. |
| **2. Synchronous & asynchronous creation** | `create_figure` | Validates inputs and renders a new figure synchronously (`workspace`, `main_tree`, optional companion trees, `options`, `clades`). | `figure_id`, `revision: 1`, MCP `resource_link` URI to `figure.svg`, normalized `spec`, and layout QA `report` (`topology_unchanged`, `remaining_collisions`). |
| | `start_figure_job` | Starts long multi-locus cloudogram or all-node dense chronogram renders asynchronously (`{ figure_spec, idempotency_key }`). | `job_id`, queue state (`queued` / `running`), and polling metadata. |
| | `get_figure_job` | Queries progress or completion of an asynchronous render job (`{ job_id }`). | Live stage progress, elapsed time, and committed `figure_id` upon completion. |
| | `cancel_figure_job` | Terminates a queued or running background job (`{ job_id }`) and reclaims workers cleanly. | Confirmed `cancelled` status without leaving partial revisions. |
| **3. Versioned semantic editing & history** | `inspect_figure` | Reads a committed figure (`{ figure_id, revision? }`) without rasterizing. | Active spec, layout/output dimensions, discordance counts, and stable semantic IDs (`clade_id`, `source_id`, `branch_id`). |
| | `edit_figure` | Applies an atomic list of semantic `operations[]` (`{ figure_id, base_revision, operations }`). | Committed `revision: k + 1`, updated SVG `resource_link`, and fresh collision/invariant report. |

| Lifecycle Stage | Tool Name | Purpose & Exact Inputs | Returned Output |
|---|---|---|---|
| | `update_figure` | Replaces or merges specification sections (`{ figure_id, base_revision, patch }`). | Committed `revision: k + 1` (or rolls back completely if validation fails). |
| | `list_revisions` | Lists all committed revisions (`{ figure_id }`). | Chronological revision log with timestamps and redacted parameter diffs. |
| | `restore_revision` | Restores an earlier state (`{ figure_id, base_revision, target_revision }`) without deleting history. | New head revision identical to `target_revision` plus restored SVG resource. |
| **4. Artifact export & lifecycle cleanup** | `export_figure` | Exports publication artifacts (`{ figure_id, revision?, formats? }`, where `formats` accepts `"svg"`, `"png"`, `"panels"` using `options.output.dpi`). | Export status, `safe_for_export` verification, warnings, and `resource_link` URIs for requested formats. |
| | `preview_figure` | Renders a bounded raster preview (`{ figure_id, revision? }`) when visual inspection is explicitly requested. | PNG preview image and layout QA diagnostics. |
| | `get_figure_spec` | Exports the archival JSON record (`{ figure_id, revision? }`), redacting raw tree text with SHA-256 input hashes. | Redacted `{ requested, normalized }` specification record. |
| | `delete_figure` | Deletes server-side artifacts and revision files (`{ figure_id }`) upon explicit user request. | Confirmation of storage cleanup. |

#### Five domain confirmation gates in the companion Agent Skill

When you work with an AI assistant loaded with the `phylorender-mcp` skill, the assistant follows five strict confirmation rules so it never guesses biological parameters or overrides tuned defaults silently:

1. **Outgroup confirmation gate:** Before creating a *Phylogram*, *Tanglegram* (using the left primary tree's outgroup), or *Cloudogram* (passed as `options.cloud.phyparts_outgroups` to root gene trees while preserving the species-tree root), if you have not specified the outgroup, the assistant inspects the tree and asks you to confirm the outgroup taxon or clade rather than guessing from file order. In *Time Tree*, it preserves your dated root and never asks to reroot.
2. **Sample-mismatch confirmation gate:** Before creating a *Tanglegram*, the assistant runs `compare_tree_samples`. If tip sets differ, it shows the complete `primary_only` and `comparison_only` lists and waits for you to choose `intersection` or `union` (or source-name correction on equal-count spelling mismatches).
3. **Dense chronogram label gate:** If `inspect_tree` reports `160` or more candidate internal-node age labels on a *Time Tree* and you have not set an age window, the assistant asks whether to use the clean default (labeling the older `50%` of the root-to-present span while retaining all 95% HPD bars), label all nodes (`age_minimum: 0`, under a cancellable `1,800 s` render deadline), or apply a custom age range.
4. **Default-first output and override confirmation gate:** New figures always start from PhyloRender's tuned publication defaults (Arial typography, automatic portrait spacing, `1.25×` median tip-width cap, and automatic collision-free label placement). Even if a prompt asks broadly to "beautify" a figure, the assistant will not send non-default font sizes, manual spacing, line

weights, aspect ratios, or DPI overrides without first listing the exact proposed parameter values and receiving your confirmation.

5. **Local-first privacy and fast vector-SVG gate:** Ordinary creation and revision steps return vector SVG directly without slow intermediate PNG rasterization (generating PNGs or calling `delete_figure` only when explicitly requested), and local loopback calls (`127.0.0.1:8766`) never fall back to a remote endpoint upon HTTP `429` (`SERVICE_BUSY`) or validation errors, keeping unpublished tree data on your machine.

#### S2.5 | Step-by-Step Operational Workflows and Practical Examples

##### Workflow 1: Interactive figure preparation in the Web Workbench

1. **Select the workspace:** Open the Web Workbench in your browser and click **Phylogram**, **Time Tree**, **Cloudogram & Discordance**, or **Tanglegram** in the top bar.
2. **Import and validate tree data (`Data` tab):** Upload your primary Newick or Nexus tree. Depending on the workspace, attach companion support trees (*Phylogram*), confirm the file time unit and node-age source in Step 2 (*Time Tree*), choose `intersection` or `union` if tip sets differ (*Tanglegram*), or upload your ML gene-tree collection and list multi-taxon species-tree outgroups (*Cloudogram*).
3. **Root, order, and style tip names (`Layout` & `Tip Labels` tabs):** In *Phylogram*, select an outgroup taxon and click **Reroot by Selected Outgroup** (or right-click an outgroup clade's branch on the canvas and click **Reroot at this branch**). Confirm automatic Latin binomial italics under `Tip Labels`, and optionally double-click individual tips or upload a two-column CSV/TSV/TXT/`.xlsx` table (`original_name, display_name`) to format publication labels.
4. **Add single-tier or nested clade brackets (`Clades` tab):** Click **Select Clade or Taxon**, then click any internal node or branch on the SVG canvas to open the clade dialog. Enter the taxon group name, pick a color swatch, and leave `Bracket tier` on `Automatic` (inner solid tier for tribes, outer dashed tier for enclosing subfamilies) or choose `Inner tier 1` through `Outer tier 6`. Use the top canvas toolbar (`Fit window`, `Fit width`, `Alt + scroll` zoom, and `Find taxon...`) to navigate large trees.
5. **Optimize node annotations and export (`Annotations`, `Time Scale` & `Export` tabs):** Fine-tune support masking, HPD bar scope, discordance pie mode, or ICS timescale font sizes via the `− / +` steppers. Click **Optimize Node Label Placement** at the bottom of the workbench to recompute zero-overlap label slots and leader lines across the figure (individual node labels can also be dragged on the canvas). Open `Export` to set DPI (`72–1200`, default `300`) and download an editable **SVG**, **PNG**, **Export Two-Panel Split `.zip`** bundle, or **`.json`** figure specification.

##### Workflow 2: Conversational plotting and revision rollback with an AI Agent

With the MCP server connected and the `phylorender-mcp` skill active in your AI client, provide your tree file paths and biological context in plain language. Including the outgroup (or time unit) upfront allows the agent to generate the figure in a single pass:

> **Example initial prompt (*Phylogram* with dual supports):**
> *"Using `sapindaceae_ml.treefile` as the primary ML tree (bootstrap percentages) and `sapindaceae_astral.tre` as an auxiliary support tree (ASTRAL posterior probabilities, 0–1), create a Phylogram rooted on `Acer_truncatum` and `Dipteronia_sinensis` using standard publication defaults and export an SVG."*

> **Example follow-up revision prompt (semantic editing on the committed revision):**
> *"Add inner solid brackets (tier 0) for tribes Nephelieae (MRCA of `Dimocarpus_longan` and `Litchi_chinensis`, `#2b6f6b`) and Sapindeae (MRCA of `Sapindus_mukorossi` and `Koelreuteria_paniculata`, `#b85d3b`), enclosed by an outer dashed bracket (tier 1) for subfamily Sapindoideae (`#1f2937`)."*

Behind the scenes, the agent hydrates local tree files, calls `inspect_tree` → `create_figure` (`revision: 1`) → `inspect_figure` → `edit_figure` (`base_revision: 1` → `revision: 2`), and verifies `topology_unchanged: true` and `remaining_collisions: 0` in the returned report. If you want to undo a later styling experiment, say *"Roll back the figure to revision 1"*, and the agent invokes `restore_revision` without erasing intermediate history.

##### Workflow 3: Bidirectional handoff between the Browser Workbench and AI Agents

1. **Agent → Browser (interactive visual touch-up):** Ask the AI agent to batch-build all MRCA clade brackets and multi-tree pairings from your classification table and save a complete portable `.json` figure specification (embedding the local raw tree strings into `main_tree.content` and companion `content` fields alongside the normalized `options` and `clades` from `inspect_figure` / `get_figure_spec`). Open the Web Workbench, go to `Export` → **Load Figure Spec (.json)**, inspect the live SVG canvas interactively, drag any individual leader label if desired, and click **Export SVG**.
2. **Browser → Agent (manuscript-wide style standardization):** Design and polish your main phylogeny interactively in the Web Workbench until typography, stroke weights, and subfamily/tribe color swatches match your target journal's style. Click **Export Figure Spec (.json)** and hand that JSON file to your AI agent: *"Apply the exact typography, stroke weights, and clade color palette from `fig3a_spec.json` (confirming those exact overrides) to our StarBEAST3 chronogram (`fig3b.trees`, ages in Mya) and our 86-gene cloudogram, then export SVG and 600-DPI PNG files for all figures."*

### Figure S1

**Article Title:** *PhyloRender: an interactive and agent-ready package for reproducible phylogenetic visualization*
**Journal:** *Molecular Ecology Resources* (Software Note)
**Authors:** Jian He, Jingjing Chen, Zihao He, Jiamin Xiao, and Lei Xie

**(a) Panel 1 — Basal Sapindales & early-diverging Sapindaceae**

65 tips + explicit continuation guide to crown Sapindoideae (111 tips)

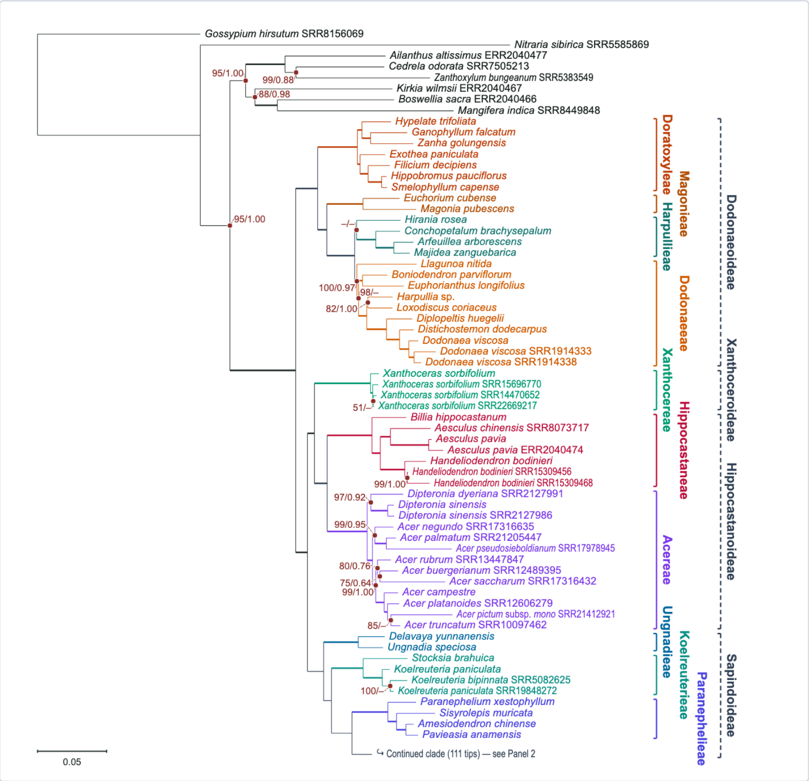


**(b) Panel 2 — Monophyletic crown Sapindoideae (111 tips)**

Inherited outer Sapindoideae bracket + inner tribe brackets & 0 collisions

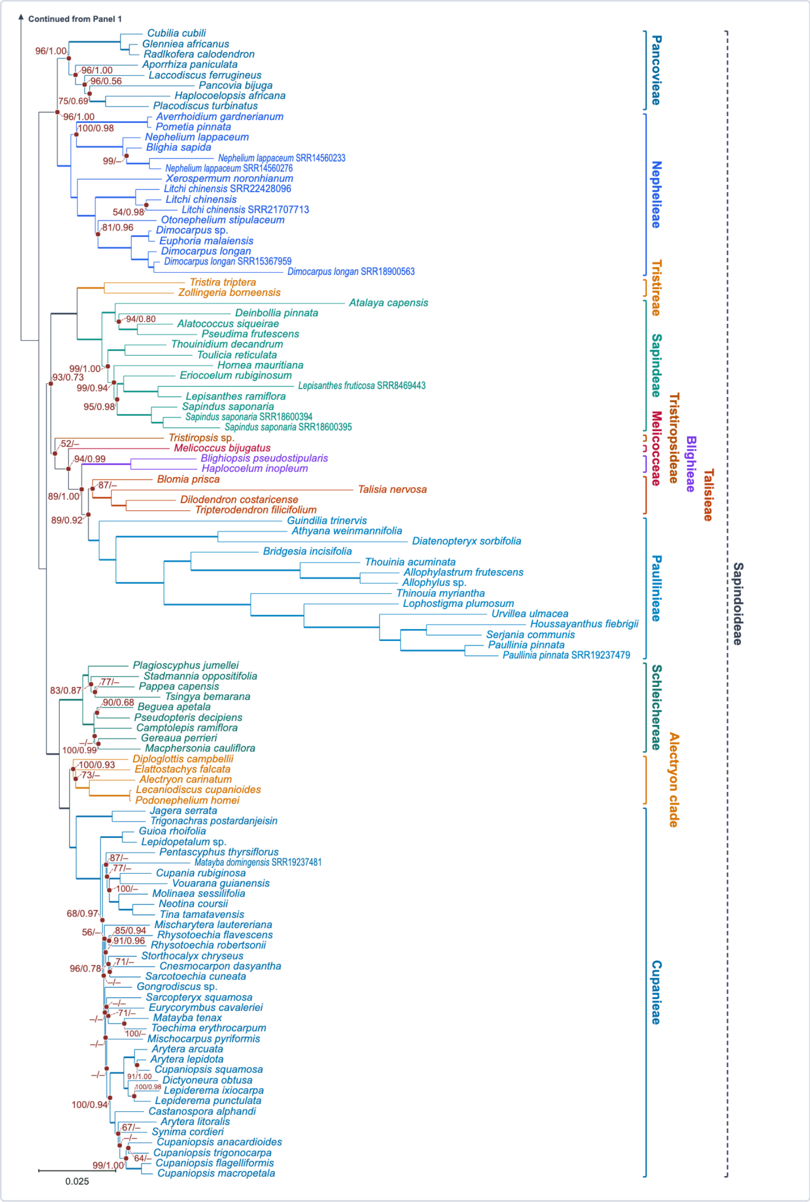

#### Legend and Description

**Figure S1. Automated two-panel split at a monophyletic clade, aligned two-tier tribe and subfamily brackets carried across panels, and dual-method support labels on the full 176-taxon Sapindaceae phylogeny.**
The 176-taxon concatenated maximum-likelihood (ML) phylogram of Sapindaceae from He et al. (2025), rooted on the outgroup *Gossypium hirsutum* (`Gossypium_hirsutum_SRR8156069`), is paired automatically with local posterior probabilities (LPP) from the corresponding 176-taxon ASTRAL coalescent species tree (Zhang et al., 2018). Following the infrafamilial classification of Buerki et al. (2021), 22 inner solid brackets mark tribes and informal clades while coloring descendant branches and tip labels, enclosed within four outer dashed brackets in slate gray marking the subfamilies Dodonaeoideae, Xanthoceroideae, Hippocastanoideae, and Sapindoideae. Each taxonomic tier is aligned into a dedicated vertical column without label overlap, and infraspecific taxon formatting (such as *Acer pictum* subsp. *mono* `SRR21412921`) italicizes Latin names while preserving raw specimen identifiers and tree topology. Branches reaching full support in both inference methods (`100%` ML bootstrap and `1.00` ASTRAL LPP) are drawn with thickened lines, values below `50%` or `0.50` are masked as `–`, and all remaining composite support labels are placed without overlap using leader lines. **(a)** Panel 1 displays the outgroups, Dodonaeoideae, Xanthoceroideae, Hippocastanoideae, early-diverging Sapindoideae lineages, and a continuation marker indicating where the crown clade is split. **(b)** Panel 2 isolates the complete 111-taxon monophyletic crown Sapindoideae clade at matching panel width while continuing the spanning outer dashed Sapindoideae subfamily bracket alongside the inner solid tribe brackets.

### Table S1

**Article Title:** *PhyloRender: an interactive and agent-ready package for reproducible phylogenetic visualization*
**Journal:** *Molecular Ecology Resources* (Software Note)
**Authors:** Jian He, Jingjing Chen, Zihao He, Jiamin Xiao, and Lei Xie

#### Table S1. Software versions, inputs, and rendering settings for the node-label comparison in Figure 2b–d

All three panels in Figure 2b–d were generated from the same 50-taxon monophyletic Sapindoideae crown radiation clade (*Plagioscyphus jumellei* to *Cupaniopsis macropetala*), extracted from the 176-taxon Sapindaceae phylogeny of He et al. (2025; Figure S1) with unmodified ML substitution branch lengths and paired `ML BS / ASTRAL LPP` node supports, and ladderized in the same orientation. Comparisons were executed with the R package `ape` (Paradis & Schliep, 2019), the Python toolkit `ETE3` (Huerta-Cepas et al., 2016), and PhyloRender. No manual label adjustments were performed in external vector graphics editors on any output.

| Panel | Software and Version | Runtime Environment | Rendering Invocation and Node-Label Strategy | Observed Behavior on Rapid Short-Branch Radiations |
|---|---|---|---|---|
| **Figure 2b** | **`ape` v5.8.1 (R package)** | R v4.4.3 (macOS arm64, Cairo PNG, 350 DPI) | `plot.phylo(ladderize(tr, right = TRUE), show.node.label = TRUE, cex = 0.56, edge.width = 1.2, label.offset = 0.001, font = 3)` with `add.scale.bar()` | Preserves true substitution branch lengths, but prints all composite `ML/ASTRAL` strings at fixed node offsets, producing severe label-on-label and label-on-branch overlaps along the short *Cupania*, *Mischarytera*, *Arytera*, and *Cupaniopsis* backbone branches. |
| **Figure 2c** | **`ETE3` v3.1.3 (Python toolkit)** | Python v3.13.9 (Qt offscreen rasterizer, 350 DPI) | `TreeStyle(show_scale=True, scale=7600, branch_vertical_margin=2.0)` with `TextFace(label, fsize=7, ftype="Arial")` at `position="float"` (`ladderize(direction=0)`). Floating placement was used rather than ETE3's default `branch-top`/`branch-bottom` mode because the latter stretches short branches with dotted spacers and alters displayed branch lengths. | Preserves true substitution branch lengths, but floating composite `ML/ASTRAL` labels cross neighboring branches and stack over one another along compressed internal edges while repeating `100/1.00` labels across full-support branches. |

| Panel | Software and Version | Runtime Environment | Rendering Invocation and Node-Label Strategy | Observed Behavior on Rapid Short-Branch Radiations |
|---|---|---|---|---|
| **Figure 2d** | **PhyloRender** | Chromium headless and Web Workbench (400 DPI export) | Phylogram workspace with automatic cross-tree branch matching pairing ML bootstrap (`%`) and ASTRAL LPP (`0–1`), thickened full-support branches, low support (`<50%` or `<0.50`) masked as `–`, and automatic candidate-position search with leader lines. | Preserves original branch lengths and topology, encodes dual full-support branches as thickened lines, and places all remaining composite support labels into nearby open space via leader lines with zero text or branch overlaps. |

### Table S2

**Article Title:** *PhyloRender: an interactive and agent-ready package for reproducible phylogenetic visualization*
**Journal:** *Molecular Ecology Resources* (Software Note)
**Authors:** Jian He, Jingjing Chen, Zihao He, Jiamin Xiao, and Lei Xie

#### Table S2. Dataset summary and layout verification across main and supplementary empirical figures

All empirical figures were generated from the published Sapindaceae phylogenomic datasets of He et al. (2025), with subfamilial and tribal brackets following Buerki et al. (2021), divergence times inferred with StarBEAST3 (Douglas et al., 2022) displayed against the official ICS chronostratigraphic chart (Cohen et al., 2013), and cloudogram overlays and discordance pies following Sanderson (2002), Smith & O'Meara (2012), and Smith et al. (2015). In the Phylogram workspace (Figures 2d, 3a, and S1), auxiliary support trees are matched to the main tree automatically by unrooted branch splits: cutting any internal branch partitions the full tip set into two complementary taxon subsets, so corresponding branches are paired reliably across inference methods even when input trees are unrooted or rooted on different outgroups. Layout verification metrics were recorded automatically by PhyloRender's export audit report for each figure.

| Figure | Workspace | Dataset and Taxon Sampling | Displayed Annotations and Clade Grouping | Label Overlaps | Data Integrity Verification |
|---|---|---|---|---|---|
| **Figure 2d** | Phylogram | 50-taxon Sapindoideae crown clade; concatenated ML tree paired with ASTRAL LPP tree | Thickened dual full-support branches; remaining composite `ML BS / ASTRAL LPP` labels placed with leader lines | `0` | Input topology, ML branch lengths, and tip identifiers unchanged |
| **Figure 3a** | Phylogram | 41-taxon Sapindaceae dataset (8 outgroups + 33 ingroup accessions); concatenated ML tree paired with ASTRAL LPP tree | Dual-method supports (thickened full-support branches and leader-line composite labels); two vertically aligned tiers of tribe and subfamily brackets across all 33 Sapindaceae ingroup accessions | `0` | Input topology, ML branch lengths, and tip identifiers unchanged |
| **Figure 3b** | Time Tree | 41-taxon Sapindaceae StarBEAST3 fossil-calibrated chronogram (`0–100 Ma`) | Internal node ages (Ma) with leader lines; 95% HPD bars; official ICS geologic timescale; two-tier tribe and subfamily brackets | `0` | Input root age, divergence times, and 95% HPD intervals preserved without support-value clutter |
| **Figure 3c** | Tanglegram | 41-taxon nuclear phylogeny vs. 29-taxon plastid phylogeny (29 shared accessions) | Paired nuclear and plastid branch supports; one-to-one shared-taxon connectors untangled by sister-clade rotation | `0` | Both nuclear and plastid topologies, branch lengths, and raw tip identifiers preserved |

| Figure | Workspace | Dataset and Taxon Sampling | Displayed Annotations and Clade Grouping | Label Overlaps | Data Integrity Verification |
|---|---|---|---|---|---|
| **Figure 4** | Cloudogram | 34-taxon dated Sapindaceae species tree with 86 complete ML gene trees | Four-category node discordance pie charts; 86 penalized-likelihood visually aligned gene-tree overlays colored by species-tree MRCA; ICS geologic timescale; two-tier clade brackets | `0` | Species-tree ages preserved; metadata explicitly record that gene-tree branch scaling represents visual alignment only |
| **Figure S1** | Phylogram (two-panel) | Full 176-taxon Sapindaceae phylogeny (8 outgroups + 168 ingroup accessions); concatenated ML tree paired with ASTRAL LPP tree | Monophyletic two-panel split isolating the 111-taxon crown Sapindoideae clade into Panel 2; cross-panel continuation of the outer Sapindoideae bracket; 22 inner tribe/clade brackets and 4 outer subfamily brackets; dual-method supports | `0` | Rooted on *Gossypium hirsutum* prior to splitting; full 176-taxon topology, branch lengths, and voucher codes preserved |

### Table S3

**Article Title:** *PhyloRender: an interactive and agent-ready package for reproducible phylogenetic visualization*
**Journal:** *Molecular Ecology Resources* (Software Note)
**Authors:** Jian He, Jingjing Chen, Zihao He, Jiamin Xiao, and Lei Xie

#### Table S3. Benchmark verification of PhyloRender's built-in gene-tree discordance module across simulated and empirical datasets

PhyloRender's built-in JavaScript discordance module implements the rooted clade comparison rules of Smith et al. (2015) directly inside the software without requiring a Java runtime or external PhyParts binaries (full mathematical formulation in `Supplementary_Methods_1.md`, Section S1.5). For each species-tree internal node, every input gene tree is first restricted to the set of taxa shared between that gene tree and the species tree. To partition conflicting gene trees into the dominant alternative clade versus all remaining alternatives when some gene trees have missing taxa, PhyloRender groups compatible conflicting partial clades using bilateral set inclusion of both the observed taxa inside each clade and the observed taxa outside it before identifying the highest-frequency alternative, eliminating any dependence on gene-tree input order. Across seven simulated and empirical benchmark suites (including empirical Sapindaceae locus sets from He et al., 2025), all four integer counts (concordant, dominant alternative, remaining alternatives, and uninformative) were compared node by node against an independent Python reference implementation of the same comparison rules.

| Experiment ID | Dataset and Experimental Design | Taxa ($N_{tips}$) | Input / Comparable Gene Trees | Evaluated Node Checks | Exact 4-Category Node Agreement | Maximum Count Discrepancy | Built-in Module Runtime (macOS arm64) |
|---|---|---|---|---|---|---|---|
| **Exp. A1** | **Exhaustive 5-taxon rooted topologies** (all 15 rooted binary species trees crossed with 16 rooted binary and multifurcating gene trees) | 5 (1 outgroup + 4 ingroup) | 240 species–gene tree pairs | **720** | **720 / 720 (`100.0%`)** | **`0`** | `244.9 ms` (total for 240 runs) |
| **Exp. A2** | **Support-cutoff boundary sweep** (`0%`, `50%`, `70%`, and `95%` strict > thresholds, including exact boundary support values `0.50`, `0.70`, and `0.95`) | 6 | 180 (× 4 cutoffs) | **16** | **16 / 16 (`100.0%`)** | **`0`** | `< 5.0 ms` |
| **Exp. A3** | **Input-order permutation invariance** (50 random permutations of mixed complete and partial-taxon conflicting gene trees) | 7 | 5 per run (× 50 shuffles) | **250** | **250 / 250 (`100.0%`)** | **`0`** | `< 5.0 ms` |

| Experiment ID | Dataset and Experimental Design | Taxa ($N_{tips}$) | Input / Comparable Gene Trees | Evaluated Node Checks | Exact 4-Category Node Agreement | Maximum Count Discrepancy | Built-in Module Runtime (macOS arm64) |
|---|---|---|---|---|---|---|---|
| **Exp. A4a** | **Empirical Sapindaceae cloudogram set** (complete-taxon ML gene trees from He et al., 2025, underlying Figure 4) | 34 | 86 / 86 comparable | **32** | **32 / 32 (100.0%)** | **0** | 4.8 ms |
| **Exp. A4b** | **Empirical angiosperm multi-outgroup set** (complete-taxon ML gene trees) | 54 | 153 / 153 comparable | **52** | **52 / 52 (100.0%)** | **0** | 19.5 ms |
| **Exp. A5a** | **Empirical Sapindaceae genome-wide locus set** (including partial-taxon gene trees with missing species from He et al., 2025) | 41 | 3,326 input / 2,784 comparable | **39** | **39 / 39 (100.0%)** | **0** | 324.5 ms |
| **Exp. A5b** | **Large-scale empirical phylogenomic locus set** (multi-outgroup rooting with extensive partial-taxon sampling) | 57 | 8,467 input / 7,304 comparable | **55** | **55 / 55 (100.0%)** | **0** | 23.44 s |

### Table S4

**Article Title:** *PhyloRender: an interactive and agent-ready package for reproducible phylogenetic visualization*
**Journal:** *Molecular Ecology Resources* (Software Note)
**Authors:** Jian He, Jingjing Chen, Zihao He, Jiamin Xiao, and Lei Xie

#### Table S4. Node-age agreement and runtime comparison between PhyloRender's built-in penalized-likelihood branch-scaling module and `treePL`

To overlay gene trees visually onto a dated species tree in the Cloudogram workspace without installing `treePL` or compiling external `ADOL-C`, `NLopt`, and `OpenMP` libraries, PhyloRender implements the rate-smoothed Poisson penalized-likelihood objective of Sanderson (2002) and Smith & O'Meara (2012) directly in JavaScript (full objective, logistic ratio parameterization, and analytical gradient equations in `Supplementary_Methods_1.md`, Section S1.5). Internal nodes whose descendant taxon sets match a clade of the dated species tree are fixed to the corresponding species-tree clade ages (`numsites = 100,000,000`, `smooth = 100`), while each remaining free node age is parameterized as a smooth logistic ratio between its parent age and the oldest fixed descendant age beneath it and optimized via a built-in L-BFGS solver. Node ages produced by PhyloRender were compared node by node against `treePL` (Smith & O'Meara, 2012) run with thorough analytical-gradient optimization (`opt = 2`, `optad = 2`, `thorough`, `moredetail`, `moredetailad`), including empirical gene-tree sets from He et al. (2025). Note that `treePL` formats output Newick branch lengths to six decimal places (`%.6f`), which introduces a baseline text-rounding floor of approximately $5 \times 10^{-7}$ age units on shared calibrated nodes.

| Experiment ID | Benchmark Suite and Calibration Design | Trees ($N_{\text{tips}}$) | Internal Nodes (Shared / Free) | Pearson *r* (Built-in vs. `treePL`) | Mean Absolute Error (MAE) | Shared Clade Max $\lVert \Delta t \rVert$ | Free Node Max $\lVert \Delta t \rVert$ | Runtime: Built-in JS vs. `treePL` (Speedup) |
|---|---|---|---|---|---|---|---|---|
| **Exp. B1** | **Simulated clocklike and rate-shifted trees** (strict molecular clock to 13× lineage rate shifts; nested fixed calibrations, root age `90 Ma`) | 20 (8 taxa) | 140 (60 / 80) | **`1.0000000000`** | $1.88 \times 10^{-6}$ Ma | $5.00 \times 10^{-7}$ Ma | $2.57 \times 10^{-5}$ Ma | `6.8 ms` vs. `325.3 ms` (**`47.6×`**) |
| **Exp. B2** | **Zero-length and micro-branch rapid radiations** (internal branches of length `0` and $10^{-7}$ substitutions/site, root age `100 Ma`) | 20 (7 taxa) | 120 (40 / 80) | **`1.0000000000`** | $5.37 \times 10^{-6}$ Ma | $2.50 \times 10^{-7}$ Ma | $2.48 \times 10^{-5}$ Ma | `3.9 ms` vs. `435.9 ms` (**`112.0×`**) |

| Experiment ID | Benchmark Suite and Calibration Design | Trees ($N_{tips}$) | Internal Nodes (Shared / Free) | Pearson *r* (Built-in vs. `treePL`) | Mean Absolute Error (MAE) | Shared Clade Max ‖ $\Delta t$ ‖ | Free Node Max ‖ $\Delta t$ ‖ | Runtime: Built-in JS vs. `treePL` (Speedup) |
|---|---|---|---|---|---|---|---|---|
| **Exp. B3** | **Cross-magnitude root-age scale invariance** (4 root-age scales spanning 10 orders of magnitude: `0.005`, `1.0`, `85.0`, and $8.5 \times 10^{7}$) | 4 scales (6 taxa) | 20 normalized nodes | **`1.0000000000`** | $< 2.0 \times 10^{-8}$ (relative) | $< 1.0 \times 10^{-12}$ (relative) | $6.00 \times 10^{-8}$ (relative to root age) | `< 2.0 ms` total |
| **Exp. B4** | **Empirical Sapindaceae cloudogram set** (He et al., 2025 dataset underlying Figure 4; root age `97.5034 Ma`) | 86 (34 taxa) | 2,838 (2,151 / 687) | **`1.0000000000`** | $3.43 \times 10^{-6}$ Ma | $6.92 \times 10^{-7}$ Ma | $5.64 \times 10^{-4}$ Ma | `0.15 s` vs. `4.46 s` (**`30.2×`**) |
| **Exp. B5a** | **Empirical 54-taxon complete ML gene-tree set** (multi-outgroup rooted angiosperm dataset; root age `58.0006 Ma`) | 153 (54 taxa) | 8,109 (3,478 / 4,631) | **`0.9999999999`** | $2.60 \times 10^{-5}$ Ma | $9.49 \times 10^{-7}$ Ma | $4.17 \times 10^{-3}$ Ma | `1.16 s` vs. `30.42 s` (**`26.2×`**) |
| **Exp. B5b** | **Empirical 57-taxon complete ML cloud subset** (complete-taxon subset extracted from the 8,467-gene set; root age `71.9994 Ma`) | 112 (57 taxa) | 6,272 (2,867 / 3,405) | **`0.9999989633`** | $1.63 \times 10^{-3}$ Ma | $6.25 \times 10^{-7}$ Ma | $< 5.0 \times 10^{-3}$ Ma on `110 / 112` trees (`0.54 Ma` on 2 flat-likelihood short-branch trees where C++ terminates early) | `0.85 s` vs. `117.12 s` (**`138.2×`**) |
